# Transcriptional re-wiring by mutation of the yeast Hsf1 oligomerization domain

**DOI:** 10.1101/2020.05.23.112250

**Authors:** Elizabeth A. Morton, Michael W. Dorrity, Wei Zhou, Stanley Fields, Christine Queitsch

## Abstract

Response to heat stress is mediated by heat shock transcription factors (HSFs), which possess conserved DNA-binding and oligomerization domains. The oligomerization domain is required for HSF1 to transition under heat stress from a monomer to a homotrimer, which alters DNA-binding specificity and affinity. Sequence variation in the oligomerization domain affects HSF1 activity, although this link is poorly understood. We performed a deep mutational scan of >400,000 variants of the oligomerization domain of *Saccharomyces cerevisiae* Hsf1 and measured fitness under stress and non-stress conditions. We identify mutations that confer temperature-specific phenotypes; some exceptional Hsf1variants lead to enhanced growth under heat stress and changes to *in vivo* DNA-binding and transcriptional programs. The link between Hsf1 oligomerization and DNA-binding domain is evolutionarily conserved, with co-evolving residues between these domains found among fungi. Mutation of transcription factor oligomerization domains may represent a path toward re-wiring transcriptional programs without mutation of DNA-binding domains.

## Introduction

The heat shock transcription factor (HSF1) is the central regulator of heat shock-inducible gene expression and the cytoplasmic unfolded protein response in eukaryotes. HSF1 has two highly conserved domains, mediating DNA-binding and oligomerization.^1, 2^ In standard models of HSF1 regulation, the protein is predominantly monomeric and inactive in the absence of stress. Upon stress exposure, HSF1 becomes post-translationally modified and trimerizes into an active state that upregulates heat shock target genes.

Despite half a century of study, the mechanisms by which HSF1 functions in the cell are not well characterized beyond this simple model. HSF1 is essential in many model organisms, including *C. elegans*, *D. melanogaster*, *S. pombe* and *S. cerevisiae*.^3–7^ Beyond regulating heat shock, HSF1 contributes to development and fertility,^3–5, 8, 9^ aging,^10^ cancer,^11–14^ innate immunity,^15, 16^ and metabolism.^17–20^ Consistent with these multiple roles, the transcriptional programs mediated by HSF1 differ in stress and non-stress conditions.^12, 21, 22^ The HSF1 DNA-binding domain is remarkably conserved, showing few changes from yeast to humans. Though initial studies of *S. cerevisiae* Hsf1 suggested it is constitutively trimerized and bound to DNA,^7^ more recent studies indicate that Hsf1 substantially increases its target occupancy upon heat shock, similar to higher eukaryotes in its temperature-dependent regulation of DNA binding.^23, 24^

Beyond regulating homotrimerization, the HSF1 oligomerization domain mediates other inter- and intramolecular interactions.^25–31^ Others have proposed that oligomerization of HSF1 can affect specificity of DNA binding,^28, 32^ and in yeast, oligomerization-defective phenotypes can be suppressed by mutation in the DNA-binding domain.^28^ The oligomerization domain of HSF1 is characterized by hydrophobic residues with heptad periodicity. Although this domain is commonly depicted as a single continuous domain, a region in the middle deviates from the ideal hydrophobic heptad repeat, although it maintains the periodicity of the flanking N- and C-terminal regions. Interruption of the hydrophobic pattern by this region, which we refer to as the HR-spacer, initially suggested that the oligomerization domain contains two distinct α-helices (HR-A and HR-B).^33, 34^ *In vitro* studies of Hsf1 protein have shown that HR-A and HR-B show differences in conformational change upon heat shock.^26^

We present here a deep mutational scan of a region of the oligomerization domain of *S. cerevisiae* Hsf1 that includes the C-terminus of HR-A, the HR-spacer, and most of HR-B. This comprehensive sequence–function map of more than 400,000 variants allowed us to expand on the role of this domain in DNA binding. Mutation of the oligomerization domain alone is sufficient to alter both target binding and the downstream composition of its transcriptional program. We identify a small subset of sites that can be mutated to confer exceptional stress resistance, with the stress-resistant mutants showing a loss of temperature-dependent repression of a subset of genes within the Hsf1 transcriptional program. These results suggest an unexpected evolutionary path to tailor the activity of a transcription factor for different environments by changing its oligomerization domain to regulate DNA binding rather than changing the DNA-binding domain itself.

## Results

### Natural variation in the HR-A/B oligomerization domain affects fitness in a temperature-specific fashion

In the consensus model of Hsf1 regulation, Hsf1 trimerizes upon heat stress and binds to heat shock elements (HSEs) to promote target gene expression (Fig. 1A). *S. cerevisiae* Hsf1 (schematic in Fig. 1B) is essential at both basal (30°C) and heat stress (37°C) temperatures, with increasing temperature accompanied by shifts in oligomeric state and changes in target specificity.^23, 35^ The oligomerization domain of the yeast protein is comprised of HR-A (heptads 1-5, residues 344-378); HR-spacer (heptads 6-7, residues 379-392); and HR-B (heptads 8-9, residues 393-403) (Fig. 1E).

**Figure 1.**
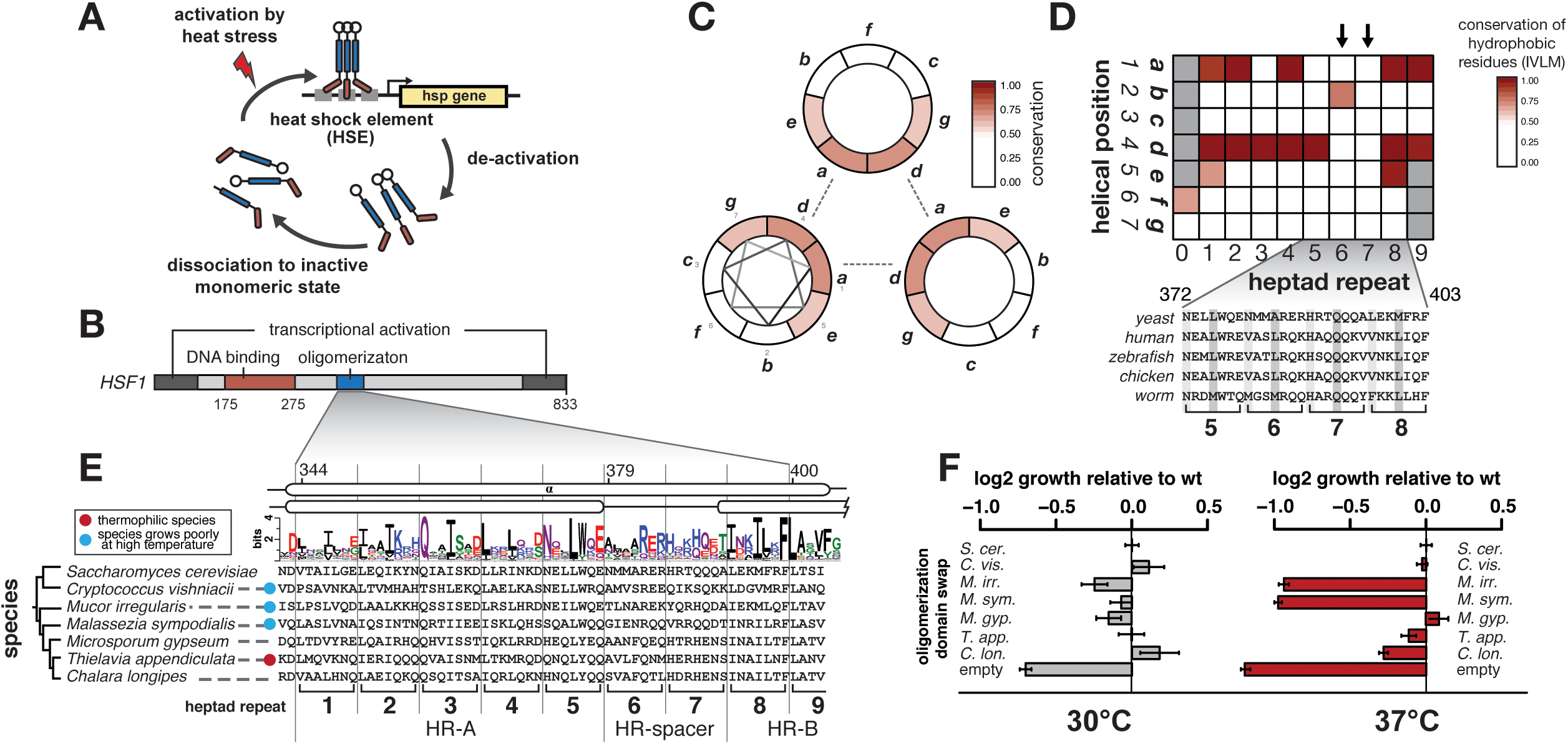
The Hsf1 oligomerization domain contributes to temperature-specific fitness and exhibits a conserved interruption of hydrophobic periodicity. (A) The standard model of induced Hsf1 activity involves conversion from an inactive monomer to a hyper-phosphorylated trimer upon stress exposure. Trimer binding to heat shock elements (HSEs) leads to upregulation of stress response genes such as heat shock proteins (HSP). The cycle is attenuated through multiple mechanisms, including interaction with heat shock proteins and acetylation.^75^ (B) The *S. cerevisiae* Hsf1 protein contains N- and C-terminal activation domains along with highly conserved DNA binding and oligomerization domains. (C) The oligomerization domain of Hsf1 has an α-helical structure (one turn = 3.6 residues). A helical-wheel representation in which each position (labeled *abcdefg*) in the helix is shown, with dotted lines indicating the predicted hydrophobic interactions in the trimer between residues *a* and *d*. Color indicates the degree of mean conservation of sites in the oligomerization domain grouped by helix position among 1229 sequenced fungal species. (D) The conservation of hydrophobic residues (Ile, Val, Leu, Met) in fungal species is broken down by helical position (*a-g*) and by heptad (1-9), showing conservation of hydrophobicity at residues *a* and *d* with the exception of heptads 6 and 7. The sequence of this equivalent region in HSFs from human, zebrafish, chicken, and *C. elegans* is shown below. (E) Six fungal species with Hsf1 oligomerization domains of either high predicted trimer propensity (red dot) or low predicted trimer propensity (blue dot) are shown below a sequence logo plot of this domain generated from fungal species. Two different predictions of α-helical regions are shown above: separation of the domain into two helixes (HR-A and HR-B)^33^ or one continuous α-helix. (F) Amino acids 342-403 of the *S. cerevisiae* Hsf1 were replaced with the oligomerization domain of one of the six fungal species shown in (E).Endogenous *S. cerevisiae* HSF1 was repressed and rescued with one of these chimeric sequences, wild-type *S. cerevisiae* HSF1, or an empty vector control plasmid. Growth at 30°C or 37°C was assayed in a plate reader overnight. Maximum slope of the resulting growth curves relative to the wild-type rescue is presented (error bars represent standard error of the mean for eight technical replicate wells).

To better understand how sequence variation in the oligomerization domain contributes to Hsf1 function, we first sought to identify patterns of sequence conservation among natural oligomerization domain variants, and to test the effects of natural oligomerization variants via domain swap experiments. We analyzed conservation of residues according to their predicted helix position in the triple-stranded coiled-coil trimer model. In the seven-residue periodicity (conventionally labeled *a* through *g*), the core *a* and *d* positions form the hydrophobic face that mediates coiled-coil interactions of the trimer.^33, 36, 37^ Like others,^33, 38, 39^ we observed stronger conservation of these core positions compared to other helix positions (Fig. 1C). As exceptions, some species show variation even at core positions, and nearly all species show a break in conserved hydrophobic amino acids at these sites within the HR-spacer (Fig. 1D, S1A). The lack of hydrophobic residues in core positions of the HR-spacer and conservation of hydrophobics outside of the core positions in HR-B suggest that these regions may encode functions beyond forming the canonical coiled-coil required for survival under heat stress.

To test if variation in this domain alone could alter the temperature-specific function of Hsf1, we replaced the *S. cerevisiae* oligomerization domain (residues 342-403) with the oligomerization domains of six fungi predicted to vary in trimerization propensity (Fig. 1E, S1B). The chimeric Hsf1 proteins were tested at 30°C and 37°C for their ability to mediate *S. cerevisiae* growth in a yeast background in which the endogenous *HSF1* was regulated to allow repression by a tetracycline derivative, anhydrotetracyclin (ATc). Substitution of the *S. cerevisiae* oligomerization domain with one from another species altered growth rate in a temperature-specific manner. At basal temperature, all oligomerization domains rescued *S. cerevisiae* Hsf1 function to at least some extent (Fig. 1F). At high temperature, oligomerization domains from three species that natively grow poorly at this temperature resulted in two (*M. irregularis* and *M. sympodialis*)^40, 41^ that caused poor growth of *S. cerevisiae*, and another (*C. vishniacii*)^42^ that fully rescued (Fig. 1F). The oligomerization domain of the thermophile *T. appendiculata*^43^ did not confer enhanced growth of *S. cerevisiae* under temperature stress (Fig. 1F). These results suggest that Hsf1 temperature sensitivity does not depend solely on the oligomerization domain.

Phylogenetic distance of the substituted domains from the *S. cerevisiae* domain does not explain differences in phenotype (Fig. 1E), suggesting that specific patterns of sequence variation in the oligomerization domain rather than predicted trimerization propensity influence temperature-dependent Hsf1 function.

### Deep mutational scanning of the Hsf1 HR-A/B domain

Because variation in the Hsf1 oligomerization domain influences temperature-dependent fitness, we sought to analyze a region of this domain by deep mutational scanning, an approach to comprehensively interrogate the functional consequences of amino acid substitutions at each position of a protein. We targeted a 36 amino acid region of the oligomerization domain (residues 365-400) spanning heptads 4 through 8, a region that includes 14 residues of HR-A, the intervening 14 residues of the HR-spacer, and 8 residues of HR-B (Fig. 2C).

**Figure 2.**
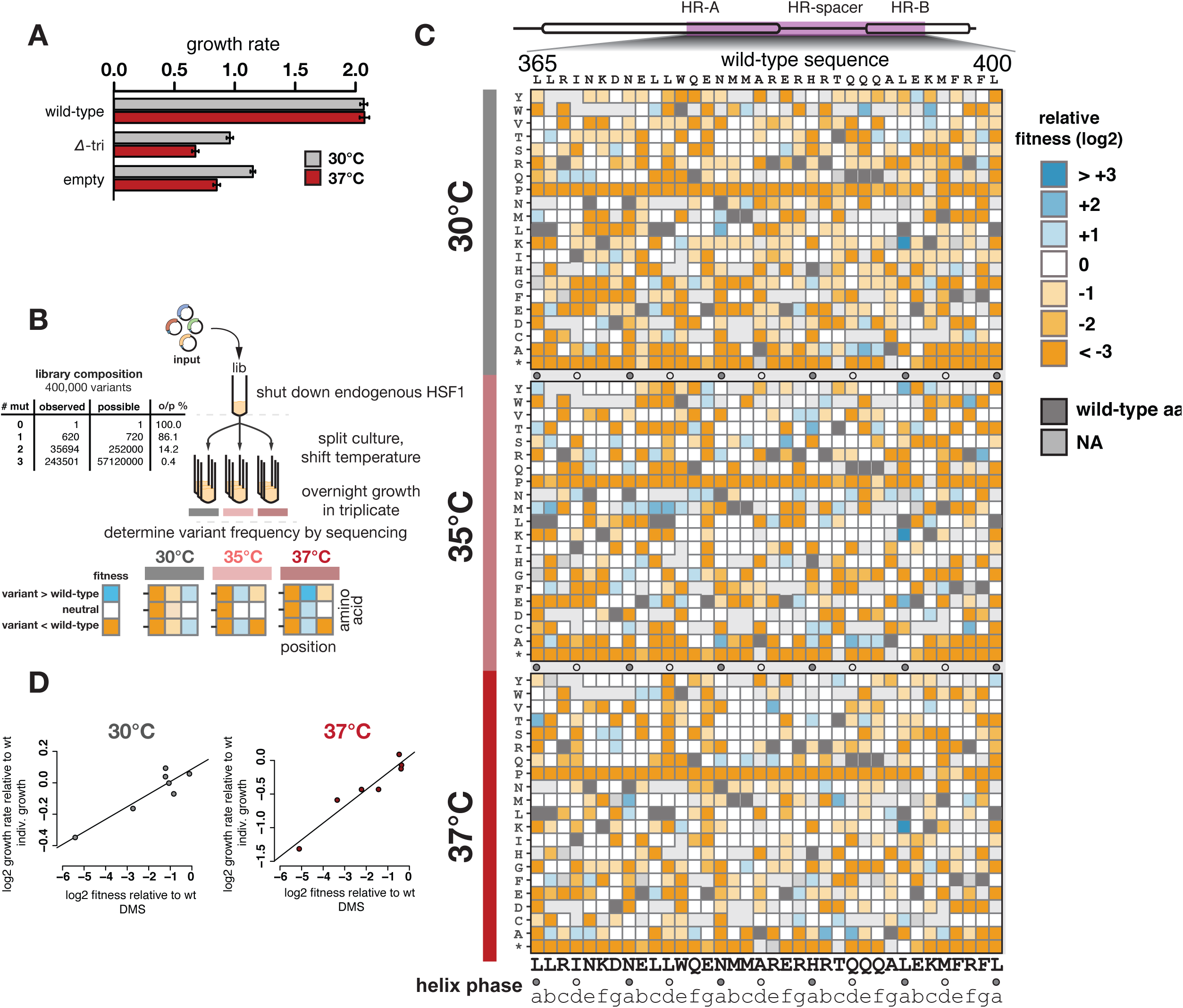
A screen of over 400,000 variants in the Hsf1 oligomerization domain reveals positional fitness effects under basal and heat stress conditions. (A) Yeast endogenous HSF1 was repressed with anhydrotetracycline (ATc) and rescued with plasmid-based expression of either a wild-type *HSF1*, an *HSF1* clone with the oligomerization domain deleted (residues 342-403 deleted), or an empty vector control plasmid. Growth of these strains was assayed overnight in a plate reader held at either 30°C or 37°C. Values on the y-axis represent maximum slope of the resulting growth curves. (B) A deep mutational scan (DMS) was conducted using a library of 400,000 variants in residues 365-400 of the oligomerization domain of a plasmid-based *S. cerevisiae HSF1*. This library contained 86% of the possible single residue changes and 14% of all possible double residue changes. The library was transformed into an *S. cerevisiae* strain with tet-off *HSF1* so that endogenous HSF1 could be repressed with ATc. The culture was split into three replicates at each of three temperatures (30°C for 14.5 hours, 35°C for 14.5 hours, or 37°C for 17.5 hours), after which plasmid libraries were extracted and sequenced for frequency. A simulated result output format is shown in (B), wherein a decrease in variant *HSF1* relative to wild type (orange) or an increase in variant *HSF1* frequency relative to wild type (blue) is presented in grid form of residue position by amino acid change, for each of the three temperatures. (C) The relative fitness scores of yeast expressing each single mutation variant in the DMS library are presented as a heat map for growth at 30°C, 35°C, and 37°C. Relative fitness determined as variant(log2(37/input)) – wild-type(log2(37/input)), where the input is the composition of the library pre-yeast transformation. Wild-type amino acid sequence is show above and below the plots, with circles marking residues at helix phase positions *a* and *d*. The region of the oligomerization domain covered in the mutagenesis is diagramed above (highlighted in purple). Variant L393K showed an enrichment score of >6.6 for all temperatures, making it more likely a sequencing error than a true enrichment. Indeed, L393K failed to show a growth advantage in validation studies (data not shown). The L393K mutation was therefore removed from subsequent analyses. (D) Seven additional *HSF1* single mutants spanning a range of fitness effects were chosen from the library and independently cloned (variants A382P, L375S, I368R, Q389K, N379I, I368L, A392N). These variants were assessed for growth in a plate reader assay as in (A). Their growth relative to wild type is plotted here against their fitness scores in the DMS.

A library of *HSF1* variants was created by doped oligo synthesis, with a 2.5% chance of mutant base incorporation at any given position. The doped oligos were inserted into a low-copy plasmid bearing the rest of yeast *HSF1,* and the library was transformed into yeast. Given Hsf1 essentiality (Fig. 2A), we used the strain allowing ATc repression of *HSF1* (Fig. 2B, see Methods), paired with complementation by a plasmid-encoded *HSF1* variant. Endogenous *HSF1* expression was permitted during transformation and expansion of the library to reduce selective pressure before we carried out the mutational screen. Some selection occurred before endogenous *HSF1* repression, attributable to putative dominant negative mutants, identified as early drop-outs (Fig. S2). Samples of the plasmid library before and immediately after transformation, as well as following overnight growth, were collected and included in downstream sequencing.

We carried out competitive fitness selections at three temperatures (30°C, 35°C and 37°C). The library culture was grown for 4 hours at 30°C in the presence of ATc to repress endogenous Hsf1 (Fig. 2B), and then split into three replicate cultures grown at each of the three temperatures for 14 hours (30°C and 35°C) or 17 hours (37°C) (Fig. 2B). Variant frequency was determined by high-throughput sequencing of the plasmids isolated from these output cultures and compared to the frequencies in the initial plasmid library, generating a fitness score. Variants that caused a fitness defect over the growth interval decreased in frequency compared to wild type, while those with a fitness benefit increased. Wild-type sequence made up 21.1% of the library. The yeast library contained 465,531 variants with at least 5 input reads, including 620 of 720 possible single amino acid changes (Fig. 2B). Correlation in variant scores was high between the three replicates at each temperature (Table S1). As expected, variants with nonsense mutations led to severe growth defects (Fig. 2C) and those with synonymous codons formed a distribution around zero (wild-type fitness) (Fig. S3). Mutations to proline were largely as detrimental as stop codons at all temperatures (Fig. 2C), consistent with an α-helical structure necessary for Hsf1 function at both high and low temperatures. We confirmed these fitness results by individually testing eight strains with variants that conferred a range of growth phenotypes. Growth rates of these strains relative to a strain bearing wild-type Hsf1 correlated with deep mutational scanning scores (Fig. 2D).

### Helical structure is a strong signal in the deep mutational scanning data

We compared the mutational scanning data to the predicted α-helical structure of the oligomerization domain (Fig. 3A and B). Substitutions at the *a* and *d* positions, particularly to small or charged residues that cannot mediate hydrophobic interactions, decreased fitness (Fig. 3A and B) at both 30°C and 37°C. This effect was not observed at the other positions (Fig. S4), with the exception of position *c*, which displayed sensitivity at 30°C to small or charged residues (Fig. 3B). The effect of helix position was also evident in double mutants: variants with combined mutations at *a* and *d* positions performed worse than those at random pairs of helix positions (Fig. S5A, S5B). We used double mutants to calculate pairwise epistasis scores between positions within the oligomerization domain. Sites enriched for epistatic interactions differed between the 30°C and 37°C conditions (Fig. S5C, S5D), supporting the idea of different conformational constraints in Hsf1 upon heat stress. The observation that sites contributing to trimeric structure are similarly sensitive to mutation at 30°C and 37°C suggests that the difference in Hsf1 regulation at these temperatures may be due to more subtle changes in the degree of trimerization, or to a mechanism other than increased trimerization, such as altered intramolecular or intermolecular interactions.

**Figure 3.**
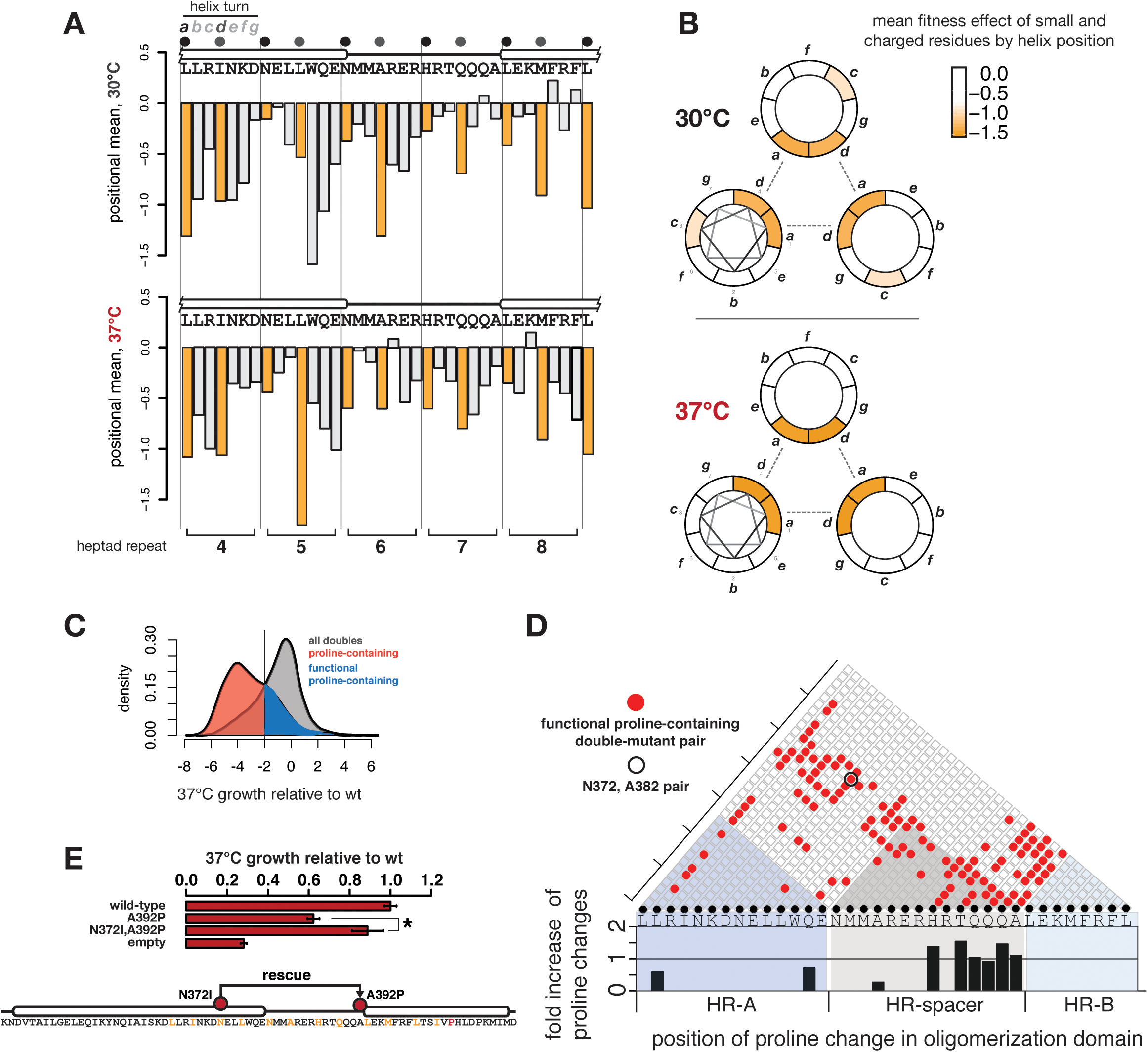
Positional effects of Hsf1 variants reveal differing mutational sensitivities for oligomerization domain in the presence and absence of stress. (A) Barplots showing the mean (log2) of fitness scores of all single amino acid changes at each position in the mutagenized region of oligomerization domain under basal (top) and heat-stress (bottom) conditions. Positions in the helical core (*a* and *d*) are colored in orange, and helical positions are indicated above. Boundaries of heptad repeats are shown with vertical lines. (B) Helix loop plots showing the mean effect of single amino changes to small or charged residues (K,R,D,E,S,G), grouped by each of the seven helix positions *a-g*. More deleterious effects are shown in shades of orange. (C) Density histograms show the distribution of 37°C fitness scores for all double mutants (gray) or double mutants that contain at least one proline (red). While the proline-containing distribution is strongly shifted in the negative direction, consistent with their unanimously deleterious effect, a fraction of these double mutants are less deleterious, and their overlap with the non-proline-containing distribution is shown in blue. (D) A pair-wise mutational plot indicates the positions of the pairs of mutations in the less-deleterious group of proline-containing double mutants (blue population in panel C). Above the wild-type sequence, each box represents a possible site pair; red circles indicate pairs found in the set of less-deleterious proline doubles. While this triangular plot does not indicate which of the two sites contributed the deleterious proline, the barplot below indicates the relative frequency of prolines at each individual site in the less-deleterious mutant set. Despite the diversity of positions among the site pairs, the contributing prolines appear highly enriched within the HR-spacer. The absence of other sites containing proline (no bar) indicates that variants at these sites were completely absent in the output population. (E) Growth rate experiments (as in Fig. 2A) independently confirm a proline double mutant identified in the screen whose deleterious effects could be mitigated by a second mutation in HR-A. Growth relative to wild type is compared between strains expressing the proline single mutant (A392P), the double mutant (A392P, N372I), and a negative control with no Hsf1 rescue (empty). A schematic of the location of each mutated position is shown below, with residues at *a* and *d* helix positions marked in orange. (error bars represent standard error of the mean for eight technical replicate wells, * t-test p-value <0.05)

We examined the effect of proline mutations specifically in the HR-spacer. Although all single mutations to proline in this region were detrimental, consistent with a required α-helix, prolines at some positions were less deleterious in the presence of a second mutation (Fig. 3C). Most functional proline-containing double mutants have at least one residue located in the HR-spacer, heptad 7 (Fig. 3D). We validated the ability of a substitution to a canonical hydrophobic residue in HR-A (N372I, position *a*) to rescue a proline substitution (A392P) in HR-spacer. Although A392P resulted in a 40% growth defect relative to wildtype, addition of N372I eliminated this growth defect almost entirely (Fig. 3E). The observation that the effects of proline mutations can be reversed more readily in the HR-spacer than elsewhere suggests that the requirement for helical structure is relaxed in this region of the oligomerization domain, consistent with distinct functions of this domain beyond coiled-coil formation.

### Exceptional Hsf1 variants increase fitness under temperature stress

A small class of variants led to a growth benefit compared to wild type at 37°C (Fig. 4A). We regenerated seven of these variants, including six double mutants (Fig. 4C), and assayed their effect on growth rate. The individual variants conferred growth rates up to 20% higher than wild type at 37°C, but less so at 30°C. One variant did not reproducibly confer the increased growth rate (Fig. S6A). We selected two of the variants, A382C and the double mutant M380K, M381I, for further characterization and found they also conferred enhanced growth under a different proteotoxic stress, exposure to ethanol (Fig. S7). We engineered a subset of the variants into the endogenous *HSF1* locus but observed no significant growth rate increase (Fig. S6B), likely a result of the better growth of yeast expressing *HSF1* alleles from the chromosome than from plasmids in the tet-off background.

**Figure 4.**
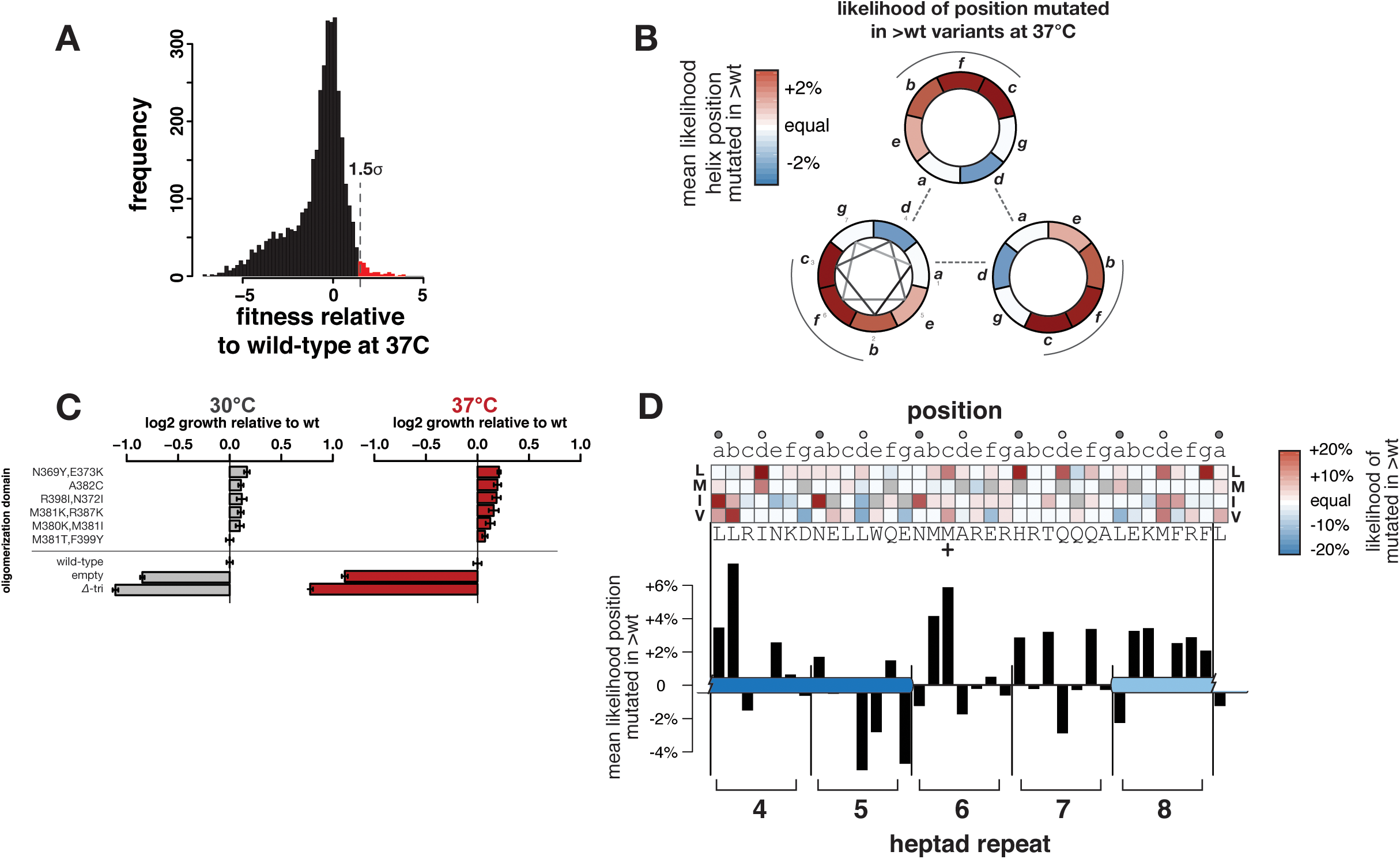
Exceptional Hsf1 variants show increased fitness under stress. (A) The relative fitness scores at 37°C for all variants are presented as a histogram. Exceptional variants were identified as those with fitness scores beyond the indicated 1.5 standard deviation cutoffs (red bars) in terms of fitness relative to wild type. (B) Variants with fitness benefits are enriched at outer helix face positions. Helix position plots reveal the frequency of mutation in this subset of variants, grouped by helix position; red shading represents an enrichment in mutation at that helix position and blue represents a depletion. (C) A set of variants chosen for their growth rate relative to wild-type in the DMS were individually validated (as in Fig. 1F) at basal (left) and heat-shock (right) temperatures. (D) Among the subset of exceptional variants, enrichment for specific substitutions (hydrophobic residues I,V,L,M, y-axis) at each position in the oligomerization domain (x-axis) was analyzed and presented as a heat map (red represents enrichment, blue represents depletion) (See Fig. S10A for full heat map). Helix positions are indicated above the map. The mean enrichment per-site is shown in the barplot below; the most enriched site is at position *b* in HR-A, and the second most enriched site is at position *c* in the HR-spacer (heptad boundaries shown as vertical bars).

Several of the mutations in the six validated variants lie between residues 380-387 in heptads 6 and 7. For the top variants conferring enhanced growth (enrichment scores >1.5sd higher than mean at 37°C), we tallied the helix positions at which mutations were present. Mutations at the predicted helix interface (position *d*) were severely depleted (Fig. 4B). However, mutations at positions *b* or *c*, predicted to be facing outward from the helix, were over-represented (Fig. 4B). Over the five heptad repeats that were mutagenized, residue 366 (in the fourth heptad) and residues 380 and 381 (in the sixth heptad) contributed the strongest signal of any *b* and *c* positions (Fig. 4D, M381 marked with “+”).

Although the pattern of many of the hydrophobic changes associated with the stress-resistant phenotype is consistent with a trimerized Hsf1, notable exceptions to the *a* and *d* periodicity occurred in heptad 4 and in the HR-spacer and HR-B. The abundance and location of these variants in the HR-spacer indicate that this region contributes to regulation of stress response by Hsf1.

### Loss of temperature-dependent regulation at a small set of transcriptional targets is associated with stress resistance of exceptional Hsf1 variants

We hypothesized that some of the mutations that conferred stress-resistant phenotypes altered the expression level, regulation or identity of Hsf1 target genes. Thus, we performed RNA-sequencing on two strains with variants that improved growth at 37°C (M380K,M381I and N369Y,E373K) and two independent strains expressing wild-type *HSF1*. Samples were collected soon after exposure to heat (30°C or 37°C for 15min or 2 hours; see Methods) or just before temperature shift but after endogenous *HSF1* repression (“+ATc”).

Transcriptional profiles of the two improved-growth variants showed high similarity to each other in genes differentially expressed compared to wild type at 37°C (Fig. S8), prompting us to pool the variants for analysis. There were few differences in gene expression between variants and wild type at the 15min timepoint; however, significant differences were observed at the 2 hour timepoint at both temperatures (Fig. 5A, 5B). Unexpectedly, those genes with significant upregulation at 37°C in the variants relative to wild type overlapped those with significant downregulation at 30°C (Fig. 5B, 5E, Table S3). Thus, the primary pattern of differentially expressed genes in the variants is the loss of temperature-specific regulation and maintenance of an intermediate expression level of these genes relative to wild-type yeast (schematic drawing in Fig. 5E inset). The exceptional Hsf1 variants showed neither constitutive nor over-expression of genes induced in the canonical heat shock response, either of which could plausibly confer increased stress tolerance (Fig. S9A, S9B). Instead, the variants failed to downregulate a set of genes – largely involved in cell wall, metabolism, Golgi-associated vesicles and cytoplasmic stress granules – in a temperature-dependent manner (Fig. 5F, Table S4). The most highly differentially expressed gene between variants and wild type at 37°C, *DED1*, encodes a protein that is globally required for yeast mRNA translation initiation and localizes to the stress granule (Fig. 5B).

**Figure 5.**
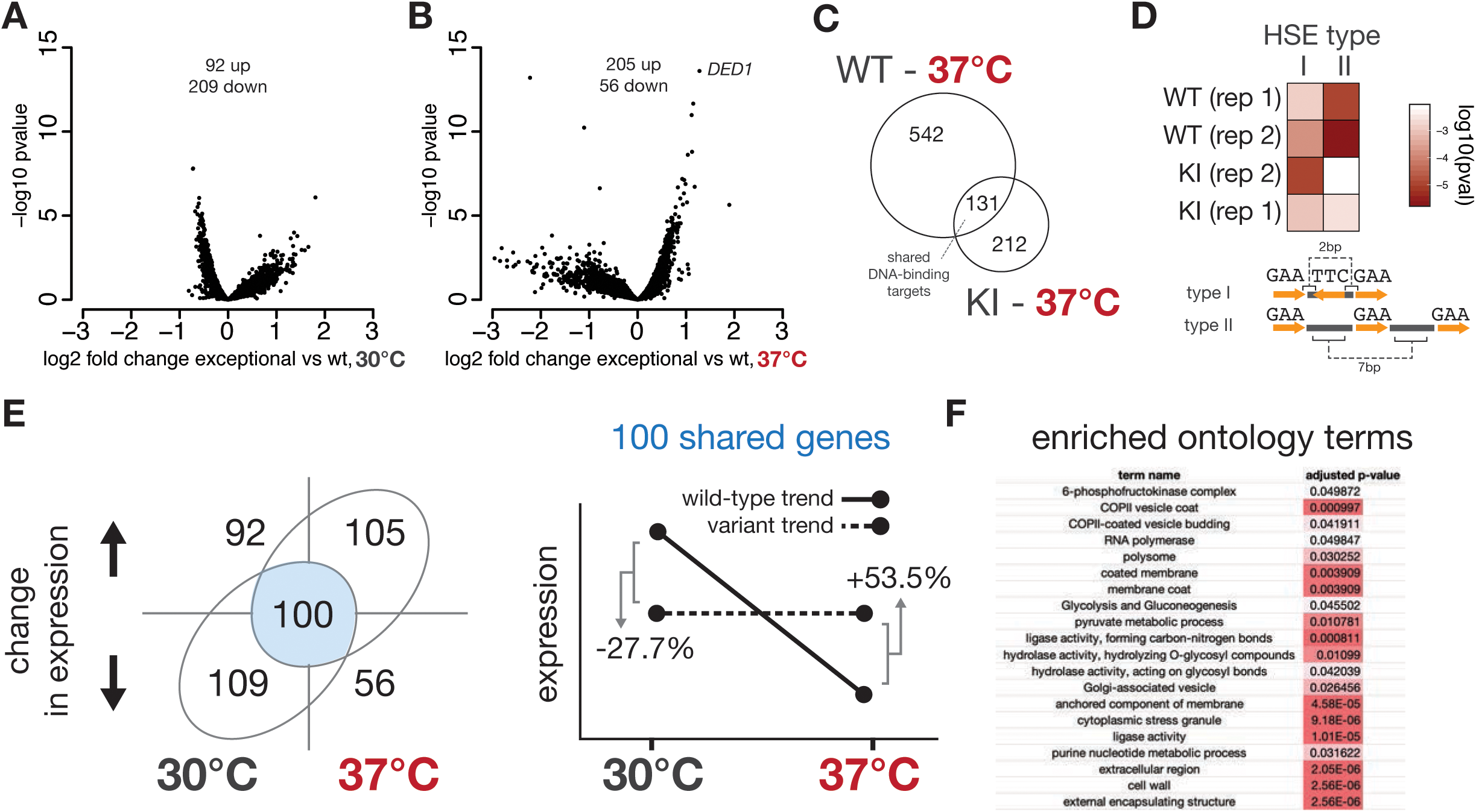
Hsf1 variants with enhanced fitness under heat stress show alterations in DNA binding and loss of temperature-dependent regulation of a set of genes. RNA-sequencing was performed on yeast with endogenous HSF1 repressed and rescued with a plasmid expressing wild type (two independent clones) or one of two HSF1 variants (N369Y, E373K and M380K, M381I, data treated as replicates in analysis). Results are presented for 120 minutes of growth. (A) For a p-value cutoff of 0.01, 92 genes were differentially upregulated in variants compared to wild type at 30°C and 209 genes were differentially downregulated. (B) At 37°C, 205 genes were differentially upregulated in the variants compared to wild type and 56 genes differentially downregulated. (C) Hsf1 binding at 37C was assessed using the transposon-insertion-based method “calling card.” The gene on either side of each high-confidence transposon insertion site was determined, tallying 673 total possible gene targets for wild type (WT) and 343 possible gene targets for variant M380K,M381I (KI), of which 131 genes were the same. (D) Motif enrichment for two different HSEs – Type I (GAAnnTTCnnGAA) and Type II (GAAnnnnnnnGAAnnnnnnnGAA) – was determined by AME (see methods) from calling card binding data for wild-type (WT) Hsf1 and the M380K,M381I variant (KI) at 37°C. Heat map values represent log10 of the p-value motif enrichment. The p-values displayed have been Bonferroni corrected for multiple sequences tested. For E-values (adjusted p-values corrected for multiple motifs) see Table S5. (E) In the RNA-seq results at the 120min timepoint, 209 genes were downregulated at 30°C in the variants with enhanced stress resistance relative to wild type. This set overlapped by 100 genes with the 205 genes upregulated in variants relative to wild type at 37°C. Most of the genes in this 100 gene list were targets significantly downregulated in wild type at 37°C compared to 30°C. The right panel of (E) shows a schematized depiction of this general trend, in which these genes are downregulated with heat stress in wild type but show more temperature-consistent expression in variants (N369Y,E373K and M380K,M381I). The average percent difference in expression level for these genes in wild type vs. variants is overlaid on the schematic drawing. (F) Gene ontology analysis revealed categories enriched in the set of 100 genes that have lost the temperature-dependent downregulation.

### Mutations in the HR-spacer domain alter Hsf1 binding targets

To understand how mutations of the Hsf1 oligomerization domain, especially in the HR-spacer, result in altered transcriptional profiles, we compared the DNA-binding patterns of strains with wild-type Hsf1 or variants. The canonical Hsf1 binding motif, the heat shock element (HSE), consists of three tandem inverted repeats of the sequence nGAAn. HSEs vary in the number and spacing of these triplet repeats among HSF1-bound genes,^12, 23, 28^ making oligomerization state a possible mechanism to differentiate among HSF1 targets.

To characterize variant Hsf1 DNA binding, we employed the calling card method.^44, 45^ In this method, *in vivo* expression of a fusion between a transposase and a transcription factor tags sites bound by the factor via transposon insertion events. Each event is molecularly barcoded so that independent binding events can be quantified. To examine changes in the sites bound under temperature stress, we applied the calling card method to wild-type Hsf1 as well as to the stress-resistant variant M380K,M381I, which falls in the HR-spacer region. Analysis of sequences flanking (150bp) transposition insertions by AME assessment identified two different types of HSEs among significantly enriched motifs at 37°C (Table S5). These two different types of HSEs (Type I, the canonical HSE, and Type II, also known as a step-type HSE^46^) exhibited different enrichment among target binding site sequences of wild-type Hsf1 and M380K,M381I (Fig. 5D and Table S5). *De novo* motif identification also found differences in significantly enriched motifs between M380K,M381I and wild type (Table S6). We conclude that a subset of the Hsf1 targeting profile remained intact in M380K,M381I, but some types of sites, particularly those with type II HSEs, differed.

We also identified the nearest gene on either side of each insertion site. Of genes associated with calling-card insertion sites, there are 131 targets in common between 37°C wild type and M380K,M381I, enriched for GO biological processes of protein folding and response to heat and enriched for HSF binding motifs (Fig. 5C, Table S7, Table S8). There are 29 genes annotated as Hsf1 targets (a significant overlap), including *HSP70 (SSA1, SSA2, YBR169C),* HSP90 complex *(HSP82, SSE1),* and co-chaperones *(YOR007C, YFL016C)* (Table S7). Wild type and M380K,M381I showed 542 genes and 212 genes unique to them, respectively (Table S7). To be inclusive in assignment of these gene lists, we included the closest gene on both sides of the insertion site. We interpret these results as evidence that mutations of the Hsf1 oligomerization domain can alter the binding specificity of Hsf1.

However, we did not observe significant overlap between the differentially expressed genes identified by RNA-seq and the genes residing near the insertion sites. For M380K,M381I at 37°C, the upregulated genes (205 genes) overlap with the calling card genes (343) by only 14 genes. For wild type at 37°C, the upregulated genes (401) overlap with the calling card genes (673) by only 43 genes, and the downregulated genes (861) overlap with the calling card genes by 124 genes. Other studies have also observed discrepancy between transcription factor binding sites and gene expression,^47–49^ attributable to such complexities of gene regulation as long-range interactions, expression changes in genes that are not direct Hsf1 targets, and the necessary but not sufficient nature of transcription factor binding.

### Natural variation in the Hsf1 oligomerization domain is consistent with its role in altering transcriptional targets

In light of our observation that yeast fitness can be modulated by mutations in the *HSF1* oligomerization domain, we examined natural variation in this domain in yeast species capable of growing under temperature stress. Using growth data for 785 industrial yeast species,^50, 51^ we identified nine thermophiles (capable of growth at 40°C or above) with available sequence data (Table S9). Residue 381, in which mutation can enhance growth under stress, shows altered amino acid preference among the nine thermophiles (Fig. 6A), with a 7-fold enrichment of hydrophobic amino acids compared to background; the consensus residue among non-thermophilic fungi was alanine (p = 0.005). As residue 381 lies within the HR-spacer (heptad 6, position *c*), a hydrophobic residue would not be expected to contribute to trimerization (Fig. 1C). To test if the HR-spacer showed expected patterns of intramolecular interactions, we examined within-domain co-evolution between heptads of the oligomerization domain across 1229 species of fungi. The trimeric coiled-coil structure predicts that heptads share co-evolving residues due to the conserved function of its quaternary structure. Consistent with this expectation, we find co-evolving sites within the same or adjacent heptad for all heptads except heptads 1 and 3 (Fig. S10B); the assumed structure of the Hsf1 trimer as an extended coiled-coil does not predict longer range contacts than these. Nevertheless, we find co-evolution of heptad 4 (HR-A) with both heptads 8 and 9 (HR-B), which extend over the HR-spacer. Co-evolution between residues distant in primary sequence can indicate physical or functional links.

**Figure 6.**
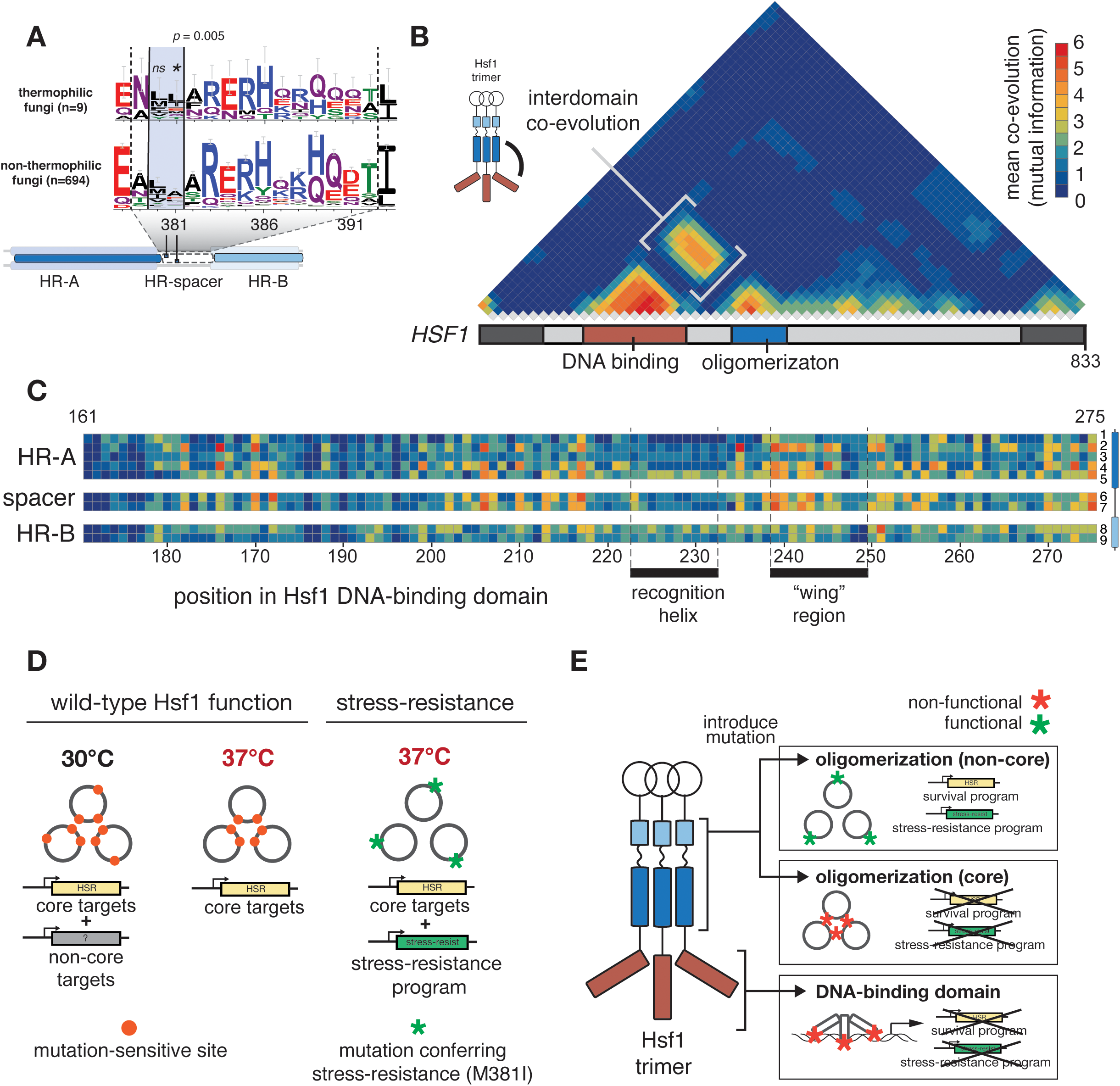
Analysis of natural variation in the oligomerization domain of Hsf1 reveals strong co-evolution between the oligomerization and DNA-binding domains. (A) Weblogo plots show the relative amino acid preferences within the HR-spacer (bound by dotted lines) for nine thermophilic fungi with a sequenced genome. Among these thermophiles, a significant (Fisher’s exact test p = 0.005) preference for hydrophobic amino acids (I, V, L, M) was found at the M381 position. The preference for hydrophobic amino acids at the M380 position was not significant (ns). The position of these sites within the oligomerization domain is shown in a schematic (HR-A in dark blue, HR-spacer with dotted lines, and HR-B in light blue). (B) Intramolecular co-evolution at the level of the entire Hsf1 protein was determined using pairwise mutational information between positions in the Hsf1, derived from analysis of Hsf1 protein sequences from 1200 fungal species. The 833^2 grid of all-by-all positions pairs was reduced to a 30^2 grid (28 aa segments of Hsf1 protein), and the mean mutual information was derived for each bin of pair-wise sites (28^2 = 784 site pairs per cell in heatmap). Binned mutual information values were smoothed with bilinear interpolation of this matrix (resolution = 1.5). Higher levels of mutual information are indicated by warmer colors. Signal near the bottom of the largest triangle indicates co-evolution between proximal residues (see triangle above DNA-binding domain). Within-domain co-evolution can also be observed in the oligomerization domain. The primary between-domain signal occurs the DNA-binding and oligomerization domains (indicated in brackets). (C) A zoom-in on all (non-binned) site pairs between the DNA-binding domain (x-axis) and heptads of the oligomerization domain (y-axis). Key regions of the DNA-binding domain are indicated with black boxes along the x-axis.^76, 77^ (D) A model schematic showing the overall pattern of mutation-sensitive sites with respect to helix position as determined by deep mutational scanning. Regardless of added temperature stress, positions in the helical core remain sensitive to mutation, while sites outside the helical core show some sensitivity in the absence of stress. The expected transcriptional targets are shown below in each case, with the heat-shock response shown in yellow gene model, and other basal targets shown in grey. In exceptional variants that activate a stress-resistance transcriptional program (green gene model), we observe an abundance of mutations on the outer helix face, away from the helical core. (E) Though exceptional variants showing stress-resistance depend on altering the wild-type transcriptional program of Hsf1, mutations in the DNA-binding domain, or in the helical core of the oligomerization domain are unlikely to retain proper targeting for the core, essential targets of Hsf1, thus leaving the outer helix face of the trimer as a viable path toward altering transcriptional targets.

Because transcription factors typically regulate target specificity via their DNA-binding domains, we would predict co-evolution between the Hsf1 DNA-binding and trimerization domains if both contribute to target specificity. Indeed, we found that mutations in the oligomerization domain co-vary with mutations in the DNA-binding domain, consistent with our experimental findings that mutating the oligomerization domain suffices to alter Hsf1 target specificity (Fig. 6B, 6C, S10C). We further analyzed co-evolution at the level of individual heptads in the oligomerization domain but failed to find specific relationships between the HR-spacer or HR-B regions and the DNA-binding domain, suggesting that interaction between these domains maybe involve the entire oligomerization domain (Fig. 6C).

## Discussion

By characterizing the effects of more than 400,000 variants of Hsf1 on yeast growth in basal and stress conditions, we found that the oligomerization domain exhibits different mutational sensitivities in basal and stress conditions, and that patterns of epistasis also vary between these conditions. These results suggest that altered interactions among individual subunits of the Hsf1 trimer contribute to temperature-specific function (Fig. 6D). Both mutational sensitivity data and patterns of epistasis point to fewer mutationally-sensitive helix positions at 37°C, consistent with a rigid, trimeric coiled-coil. In contrast, unexpected mutational sensitivity on the outer helix face and a more complex pattern of epistatic interaction at 30°C suggests that the functional Hsf1 unit, while trimeric, may depend on additional residues in the outer helix face for complete activity (Fig. 6D). Additionally, we identified exceptional variants in this domain that confer enhanced growth at 37°C, a phenotype accompanied by changes in transcriptional program and DNA targeting (Fig. 6E). Many of these variants are double mutants, such as M380K,M381I, for which the single composite mutations alone did not confer the 37°C benefit in the screen (Fig. 2C). We propose that the HR-spacer and HR-B have an activity in addition to mediating trimer formation. The exceptional variants appeared to have lost temperature-dependent regulation – either directly or through an indirect target – of a suite of genes involved in metabolic processes, transport, cell wall and cytoplasmic stress granules, likely due to the link between DNA-binding domain variation and oligomerization domain variation. In this case, the variants have resulted in cells that maintain an intermediate expression level that does not change under stress, suggesting loss of temperature-dependent regulation underlies their stress-resistant phenotype. We predict that these specific mutations would not be beneficial under other conditions, such as a fluctuating environment. The prevalence of HSF1 gene duplications, especially in plant species, allows the exploration of novel regulatory programs by introducing variation in other copies. Indeed, variable length (in residue multiples of seven) of the HR-spacer has been used to distinguish between classes of HSFs present in plants.^52^

It is possible that mutations at non-canonical positions in the HR-spacer or HR-B affect Hsf1 function through intra-molecular interaction. This model is also consistent with studies noting that HR-B may act repressively in Hsf1 functional assays.^34, 53, 54^ It is also possible that the variants that confer exceptional growth promote a different multimerization state. Hsf1 occurs in oligomeric states higher order than a trimer; *Drosophila* HSF has long been considered hexameric in its active state, with hydrophobic residues in HR-B in a register one removed from the main register possibly involved.^38^ These data also do not rule out altered interactions of other regulators of Hsf1 or disrupted post-translational modification.

While re-wiring the Hsf1-dependent stress-responsive regulatory network may be advantageous under some conditions, maintaining the core heat-stress program is necessary for survival (Fig 6D). Similar rewiring of the Hsf1 transcriptional program has been observed in cancer cells, where Hsf1-dependent regulation and translation and metabolism genes drive cell growth in a way that is distinct from heat stress-dependent regulation.^12, 55^ There are parallels between the effects seen in cancer cells and those we observe: changes to Hsf1 regulation have led to a “pro-survival” phenotype that emphasizes cell growth even in the presence of a strong stress signal.

We suggest that more subtle changes to the Hsf1 regulatory network can be made through mutations in the trimerization domain than through mutations in the DNA-binding domain itself. Hsf1 has flexibility in recognition of its DNA-binding site, with different spacings between frequently degenerate GAA motifs.^32, 46, 56^ For this reason, the potential set of binding targets for Hsf1 is large compared to transcription factors with longer sequence motifs and less variability in their arrangement. Furthermore, the requirement of a trimeric structure for high-affinity binding to DNA imposes a constraint on mutations that alter affinity for DNA; the binding affinity consequence of any single change to the DNA-binding domain is three-fold in the Hsf1 trimer.

Nevertheless, re-wiring of transcriptional networks is rampant in evolution, and altering the transcriptional targets of Hsf1 would be especially advantageous in changing how specific sets of genes are regulated by temperature. Changes to the oligomerization domain may represent a more viable path toward subtly altering transcriptional targets of Hsf1 (Fig. 6D) without overtly compromising the underlying heat shock response. The possibility that new transcriptional options are exposed by modification of the oligomerization domain has broader implications for the many ways in which species can adapt transcription factor function over evolutionary time.^57–61^

## Methods

### *Deep mutational scan of* S. cerevisiae HSF1 *oligomerization domain*

The *HSF1* gene of *S. cerevisiae* was cloned under its native promoter (664bp upstream of the start codon through 693bp downstream of the stop codon, see Table S10) into vector pRS415 (*LEU2* selection).^62^ In order to facilitate large scale library cloning, silent restriction enzyme sites were introduced on either end of the target region - AvrII (CCTAGG) at base pairs 1041-1046 and Bpu10I (GCTTAGG) at base pairs 1244-1250 – to generate plasmid pEM74. The region targeted for mutagenesis was base pairs 1093-1200 of the *HSF1* coding sequence. A doped oligo library of 168 bases was ordered (TriLink Biotechnologies) containing 30 bases of invariant flanking sequence on either side of the 108 bases of the doped target region (2.5% base misincorporation). pEM74 was digested sequentially for 7.5hr each with AvrII and Bpu10I and purified (Zymo Clean & Concentrator D4003) to generate the vector component in which to clone the doped oligo library. Flanking sequences in the doped oligo library were extended with primers to increase overlap with the digested pEM74 (Table S10, the library was amplified with primers EM244 and EM245 for 9 cycles 98°C 15sec, 60°C 15sec, 70°C 15sec). Gibson assembly (NEB #E2611S) was employed to insert the amplified library into the AvrII- and Bpu10I-digsted pEM74. The ligated variant library was cloned into DH10B cells (Invitrogen 18290015), grown in bulk, purified, and transformed using lithium acetate (see below) into tet-off *HSF1* yeast (KanMX::tetO::minpromoter::HSF1; URA3::CMV::tTA; his3-1;leu2-0;met15-0; MATa).^63^ Post-transformation, the yeast library was recovered overnight in 400mL C-Leu media, concentrated, and frozen in 25% final concentration glycerol.

The yeast library was subjected to selection and screening conditions as follows. One hundred milliliters of C-Leu media was inoculated with ∼1.2 x 10^8^ cells of frozen yeast library stock and grown at 30°C overnight. The culture was back-diluted to 100mL of OD600 0.45, at which point the remaining overnight culture was pelleted and frozen as “overnight” timepoint. The back-diluted culture was grown at 30°C for 2hr, anhydrotetracycline (ATc) (Sigma-Aldrich 37919-100MG-R) was added to 0.1µg/mL, and the culture was grown at 30°C for another 4hr. The culture was diluted to OD600 0.04 in three flasks of 100mL C-Leu + 0.1µg/mL ATc. A sample was taken as the “+ATc” timepoint. Flasks were put shaking at 200rpm at temperatures 30°C, 35°C, or 37°C and grown 14.5hr (37°C was grown 17.5hr). After growth, the 100mL cultures were spun down and pellets were frozen as post-selection timepoints.

Plasmids were extracted from pre- and post-selection libraries, as well as a sample from the frozen library stock, using the Zymoprep Yeast Plasmid Miniprep II kit (D2004). These minipreps were treated to remove genomic DNA contamination with combined Exonuclease I (NEB M0293L) and Exonuclease III (NEB M0206L) for 30 min at 37°C, followed by heat inactivation at 95°C for 2 minutes.

Each plasmid miniprep sample was amplified by Phusion (NEB M0530L) to generate libraries of the mutagenized HSF1 region flanked by cluster generating Illumina adaptors. Each sample was amplified with a different index sequence in the primers for subsequent demultiplexing (Table S10). Library amplified products were 320bp (including adaptor sequence). Following amplification, libraries were Zymo Clean & Concentrator purified (Zymo Research D4003), eluting in 15L of nuclease-free water. Libraries were quantified with Qubit high sensitivity (Invitrogen Q32854) and diluted to 2nM. Final library denaturation and dilution was performed as described (NextSeq Denature and Dilute Libraries Guide 15048776 Rev. D). Libraries were sequenced with a NextSeq High v2 300 Cycle kit (FC-404-2004), allowing full paired end coverage of the 108bp mutagenized region.

### Analysis of deep mutational scanning data

Using ENRICH software, read counts for each variant before (input) and after selection (output) were used to determine fitness of *HSF1* variants. Briefly, counts for a particular variant in the input and output libraries were normalized by their respective read depths to determine frequency in each, and a ratio of the output and input frequencies determine a variant’s functional score. Finally, enrichment scores are normalized by the enrichment of wild-type Hsf1 in each selection. Intramolecular epistasis scores (as in S5C) were calculated for all double mutants where constituent single mutants were tested; the value of epistasis is defined as the deviation of double-mutant’s fitness score from the multiplied scores of its constituent singles. Negative epistasis scores indicate that a mutation’s effect is exacerbated by the presence of a partner mutation, while positive epistasis scores indicate that a mutation’s effect is diminished by the presence of a partner mutation. Analyses of mutational frequencies (as in Fig. 4D) that include single, double and higher order mutants were completed by comparing two sets of trimerization sequences: 1. A query set ranked by fitness scores (fitness >1.5 standard deviations above mean 37°C fitness for Fig. 4D) 2. A background set with neutral fitness scores (within 0.5 standard deviations of the mean in Fig. 4D). Within each set, we calculate the frequency of each amino acid at each site to generate a frequency matrix. Enriched amino acid changes in the query set are identified by calculating the ratio of the query set’s frequency matrix to the background set’s matrix.

### Analysis of HSF1 HR-A/B oligomerization domain conservation in fungal sequences

Using fungal genomes deposited into NCBI and from the fungal sequencing project at Joint Genome Institute, we created a BLAST database of translated coding sequences and queried with the *S. cerevisiae* Hsf1 protein sequence. Full protein sequences from BLAST hits (1,229) were aligned using MUSCLE ^64^ and used to determine conservation at each site in the mutated segment. Plots showing relative conservation of each amino acid per site were generated with Weblogo.^65^ Co-evolving pairs of sites were identified using the MISTIC ^66^ tool, wherein mutual information is calculated between amino acid identities at all site pairs.

### *Cloning of* HSF1 *variants*

Point mutations identified from the DMS were introduced into pEM74 by the following method: inverse PCR was performed on plasmid pEM74 with primers that amplified outside the region of the desired nucleotide change. Gibson assembly (NEB #E2611S) was then performed with the inverse PCR and a 60-base oligo containing each desired mutation (Table S10 for inverse PCR primers and oligos, along with the list of all mutants independently cloned outside of the library). All clones were confirmed with Sanger sequencing. *HSF1* variant clones were then transformed into the tet-off *HSF1* strain^63^ by “one-step” transformation. “One-step” transformation: 1mL of saturated culture was pelleted, and 1µL of plasmid DNA (approximately 200-400ng) added to the pellet, followed by 75µL of one-step buffer [0.1M lithium acetate, 38% PEG-3350, 0.1M DTT, 0.5mg/mL carrier DNA (Roche 1467140)], after which the cells were vortexed, incubated in a 42°C water bath for one hour, and plated on C-Leu selection plates. The Δtri mutant was created by ligation (using KLD enzyme mix NEB E0554S) of an inverse PCR that omitted the HR-A/B domain (amino acids 342-403) (Table S10 for inverse PCR primers). *S. cerevisiae* HR-A/B domain (amino acids 342-403) were replaced with HR-A/B domains of other fungal species by inverse PCR and Gibson assembly with synthesized gBlocks of the domains (Table S10). The HR-A/B amino acid coordinates of the species used: *Cryptococcus vishniacii*(510-571), *Mucor irregularis* (305-366), *Malassezia sympodialis*(142-203), *Microsporumgypseum* (99-160), *Thielaviaappendiculata* (288-349), *Chalara longipes* (314-375).

### *Growth validation assays of* HSF1 *variants*

Variants identified in the DMS were validated in a plate reader growth assay. *HSF1* variants expressed under the native *S. cerevisiae* promoter on a pRS415 backbone were transformed into a background of tet-off *HSF1* (see above). Strains were cultured overnight at 30°C in C-Leu media and backdiluted in the morning to an OD600 of 0.45. Backdiluted cultures were incubated at 30°C for 2hr, after which ATc was added to a final concentration of 0.1µg/mL. The cultures were incubated for another 4hr to allow time for endogenous *HSF1* repression. Strains were backdiluted in C-Leu+0.1µg/mL ATc to an OD600 of 0.1 in a 200µL volume per well in 96-well culture plates. Eight technical replicate wells per strain per plate were run. Plates were sealed with a plate cover and put in a BioTek Synergy HI microplate reader set to 30°C or 37°C. 30°C and 37°C plates for each experiment were prepared together and run simultaneously on two different microplate readers. Plates were incubated with orbital agitation for ≥ 23 hr with 660 absorbance readings taken every 10min. Figure 4C represents eight technical replicates of one set of transformants. An independent set of transformants is presented in supplemental Figure S6A. The set of variants presented in Figure 2C was performed in tet-off *HSF1* yeast also containing an *SSA4* promoter driving GFP on a pRS413 backbone and so were cultured in C-Leu-His media with ATc.

Exposure to ethanol stress was performed by growing above plasmid-expressed *HSF1* variants in a background of tet-off *HSF1* overnight. Cultures were backdiluted to OD600 0.45, grown at 30°C for 2hr, after which ATc was added to a final concentration of 0.1µg/mL and the cultures were grown for another 4hr. Cultures were back-diluted again to OD600 0.1 in the presence of 0.1µg/mL ATc and one of the following ethanol conditions: 0%, 5%, 8%, or 10%. As above, these dilutions were grown in a BioTek Synergy HI microplate reader set to 30°C or 37°C. Four technical replicate wells were run for each variant and empty vector, and eight technical replicate wells were run for the wild type Hsf1 plasmid control.

### *Calling card analysis of* HSF1 *variants*

*HSF1* sequence with point mutations were introduced into helper plasmid of PiggyBac-based transposon system.^45^ Each helper plasmid was paired with a donor plasmid containing specific barcode within transposon. All constructs were confirmed with Sanger Sequencing. Paired donor plasmid and helper plasmid were transformed into yeast (BY4705 MATα) by high-efficiency lithium acetate transformation.^67^ After transformation, cells were collected and put on the induction plates with galactose. Cells were induced for 5 days in either 30°C or 37°C (heat shock condition) to express the HSF1-PBase. Cells were then collected, diluted back to OD600=0.45, and cultured in rich medium for 6 hours recovery (to OD600=∼1.6). Cells were put onto selection plates with 5-FOA and G418 at varying dilutions and grown 2-3 days. Cells were then collected from selection plates and genomic DNA was extracted from each sample using the Smash and Grab method as described.^68^ Four variants of *HSF1* in addition to wild-type *HSF1* were expressed for this assay and exposed to 30°C or 37°C in two replicates; all of the *HSF1* variant-expressing yeast failed to produce enough colonies for collection at 30°C, therefore only wild type had data for both 30°C and 37°C. Other variants and wild-type had two replicates per temperature, except A382P 37°C, where a single sample was collected. The library for each group were prepared through the protocol described by the calling card-seq method.^45^ The PCR products were pooled and Illumina sequenced.

The raw data included all FASTQ files from the PiggyBac transposase-only control, wild type HSF1 and the four variants: M380K,M381I (phenotype of enhanced growth at 37°C), M380I,M381K (mildly enhanced or approximately equal to wild type, Fig. S6A), Q389K (mildly impaired growth), and A382P (severely impaired growth), two replicates each. In read1, the first segment is the universal primer sequence: CGTCAATTTTACGCAGACTATCTTTCTAGGG followed by the flanking genome sequence of 39 bp. The first 8 bp of read2 is the UMI sequence used to identify unique insertion events. We first filtered sequence reads with high quality and mapped them back to the yeast genome. Then we quantified independent PiggyBac insertions based on the UMIs detected in each sample. Target genes were then assigned to insertion peaks that were within 1000 bp 5′ or 200 bp 3′ of the transcription start site for that gene.^44^

Insertions with counts above the 85th percentile were identified as “high count insertions.” We identified 300 bp windows around each high count insertion, and then merged the windows (bedops -m) to generate high insertion count sites for each replicate. We extracted the sequence from these windows and attempted to identify *de novo* motifs using MEME (Table S6).^69^ We used AME ^69^ to determine enrichment of known motifs in the target sequences. For AME, sequences (300bp: 150bp flanking each high-cut-count insertion site) were compared to a reference database of motifs.^70^ Significance values are given as reported by AME; in summary, for motif enrichment in target sequences p-values were calculated by one-tailed Fisher exact test compared to occurrence in 2-mer shuffled control sequences, with Bonferroni correction for the number of sequences input for each sample. E-values reported by AME represent p-values corrected for the expected number of motifs enriched in the test sequences given the testing of multiple motifs. (AME results Table S5).

A gene target list was generated by identifying the closest gene on either side of the high-cu-count insertion site. Both genes were included in the final gene list per sample, with duplicate gene IDs removed (Table S7). The overlap in gene lists for the two replicates of a given genotype was used as that genotype’s final gene list. Gene Ontology enrichment was performed using gprofiler (default parameters of g:SCS significance threshold multiple testing correction)^71^

### *RNA-seq of* HSF1 *variants*

RNA-seq was performed on yeast bearing *HSF1* variants transformed into tet-off *HSF1* yeast.^63^ Early timepoint RNA cultures were collected for transformants of plasmids bearing *HSF1*(N369Y, E373K), *HSF1*(M380K, M381I), or two independent clones of plasmid rescue with wild-type *HSF1* (all clones containing silent restriction sites). Clones were grown overnight at 30°C in C-Leu media and back-diluted the following morning to OD600 0.45. These back-dilutions were grown for 2hr at 30°C and then ATc was added to a final concentration of 0.1 µg/mL and the cultures were given 4hr more at 30°C. 25mL of this culture was collected for the “+ATc” sample. Cultures were diluted to OD600 0.1 in C-Leu + 0.1 µg/mL ATc in two sets of flasks, one of which was placed at 30°C and the other at 37°C. Samples of both 30°C and 37°C cultures were collected after 15min and after 2hr at their respective temperatures. Collection of each timepoint was done as follows: 50mL of the cultures was spun 4000 rcf 5 min, the pellet resuspended in the residual media and transferred to a 1.5mL tube, then spun again at 21130 rcf for 5min and the supernatant removed, immediately after which the pellet was flash frozen in liquid nitrogen.

RNA was isolated via acid-phenol extraction.^72^ RNA was DNase treated with RQ1 DNase (Promega M6101) by addition of 5µL RQ1 buffer and 5µL RQ1 DNase to each RNA sample, incubation at 37°C for 45min and then stopping the reaction with 5µL provided stop solution.

cDNA was generated with Superscript IV reverse transcriptase (Invitrogen 18090050). 5µg of RNA was combined with oligo d(T)18 primers and dNTPs according to manufacturer instructions and incubated 65°C 5min. Reactions were put on ice 2min and then combined with 4L 5X SSIV buffer, 1µL 100mM DTT, 1µL RiboLock RNase inhibitor (Thermo Fisher K1622), and 1µL SuperScript IV reverse transcriptase. The reaction incubated at 55°C for 15min followed by 80°C for 10min before being put on ice. Second strand synthesis was performed with the NEBNext Second Strand Synthesis module (NEB E6111S). 48µL of nuclease-free water was added to each tube, followed by 8µL of 10X Second Strand Synthesis reaction buffer and 4µL Second Strand Synthesis enzyme mix. The reaction incubated at 16°C for 2.5hr, after which the cDNA was purified using the Zymo Clean & Concentrator kit (Zymo Research D4003).

Libraries were generated via tagmentation of cDNA (Illumina FC-121-1030) and sequenced. Gene Ontology enrichment was performed using g:Profiler (default parameters of g:SCS significance threshold multiple testing correction) and category list for Fig. 5F condensed with REVIGO.^71, 73^

### Replacement of endogenous HSF1 with HSF1 variants

Integration of *HSF1* variants at the endogenous locus was done using Gibson assembly (NEB #E2611S) to create clones of three *HSF1* variants: A382C; M380K,M381I; and N369Y,E373K. These versions of *HSF1* had otherwise wild-type sequence and were lacking the restriction sites introduced in the cloning process of the original deep mutational screen (see above). 664bp of promoter was included upstream of *HSF1* along with 430bp of downstream sequence before a *URA3* cassette flanked by homologous recombination arms.^74^ The *HSF1* variant plus *URA3* cassette was amplified using primers with an additional 43bp of *HSF1* downstream sequence added to the 3’ end (Table S10). This PCR product was transformed with lithium acetate into yeast strain BY4741 and plated on C-Ura. Resulting recombinants were genotyped using PCR and Sanger sequencing (Table S10). Strains with *URA3* integration but wild-type *HSF1* sequence were also isolated as matched controls (Fig. S6B). Genotyped strains were then grown on 5-fluoroorotoic acid (5-FOA, 1g/L) to counter select for *URA3* and generate a line with the uracil auxotrophy. Genomic DNA was extracted (Zymo #D2002) from colonies grown on 5-FOA and Sanger sequenced to confirm presence or absence of *HSF1* mutation and *URA3* gene.

### Lithium acetate transformation protocol

One hundred milliliters of culture was grown to approximately OD 1.8. The culture was pelleted 3min 3000rpm. Supernatant was removed and cells were resuspended in 15mL LiSorb (lithium acetate, 1M sorbitol, Tris, EDTA). Cells were pelleted and resuspended in 15mL LiSorb again, and then pelleted and resuspended in 1.5mL LiSorb. The cells were split between two 1.5mL tubes and allowed to incubate with rotation at room temperature for 30min. 200µL of the cell suspension was added to four tubes, each containing 50µL of 2mg/mL salmon sperm DNA and 3µL DNA for transformation (500ng/µL DMS library or PCR product for integration(84 to 231ng/µL)). 1mL of LiPEG (lithium acetate, Tris, EDTA, 44%PEG) was added and the tubes were allowed to incubate with rotation at room temperature for 30min. 100µL DMSO was added and the tubes were incubated 42°C for 10min, after which the cells were pelleted 1min 5000rpm, the supernatant removed, and the pellet resuspended in YPD+0.5M sorbitol. The culture was allowed to recover at 30C for 1hr. For PCR product transformation, the culture was then pelleted and plated on C-Ura plates. For DMS library transformation, the cells were added to 400mL C-Leu media and allowed to grow overnight before concentrating and freezing.

## Data Availability

Expression data are available at the Gene Expression Omnibus (GEO number: pending).

## Acknowledgements

We thank Dr. Kerry Bubb for valuable assistance in data processing. This work was supported by the National Institutes of Health [Genetic Approaches to Aging Training Grant 4 T32 AG 57-39] and the National Human Genome Research Institute [Genome Sciences Training Grant 5 T32 HG 35-19] to E.A.M., and also the NSF GRFP and WRF-Hall Fellowship to M.W.D. The work was also supported by NIH grants GM114166 and 1RM1HG010461 to C.Q. and S.F.

---

**Supplementary Figure 1.**
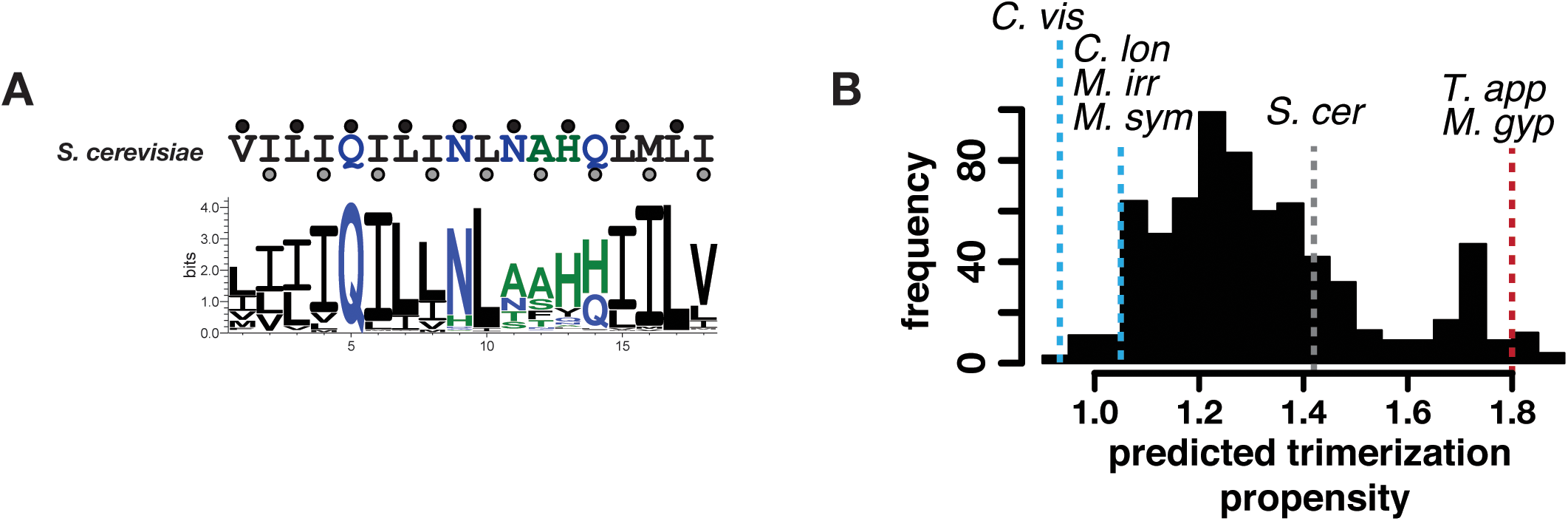
Hsf1 oligomerization domain sequences across fungi conserve hydrophobic residues at a and d positions as well as display differences in predicted trimer propensity. (A) Logo plot showing the frequency of amino acids at the helical positions a (black dot) and d (grey dot) within the Hsf1 trimerization domain. The wild-type *S. cerevisiae* sequence at these a and d positions is shown above for reference (V344 to I403). (B) Histogram showing LOGICOIL model outputs of trimerization propensity for oligomerization domains of 1229 sequenced fungi. Higher values indicate higher probability of that sequence conferring a trimeric coiled-coil. The prediction score for specific species tested in oligomerization domain swap experiments (Fig 1F) are indicated, along with the prediction for wild-type *S. cerevisiae*.

**Supplementary Figure 2.**
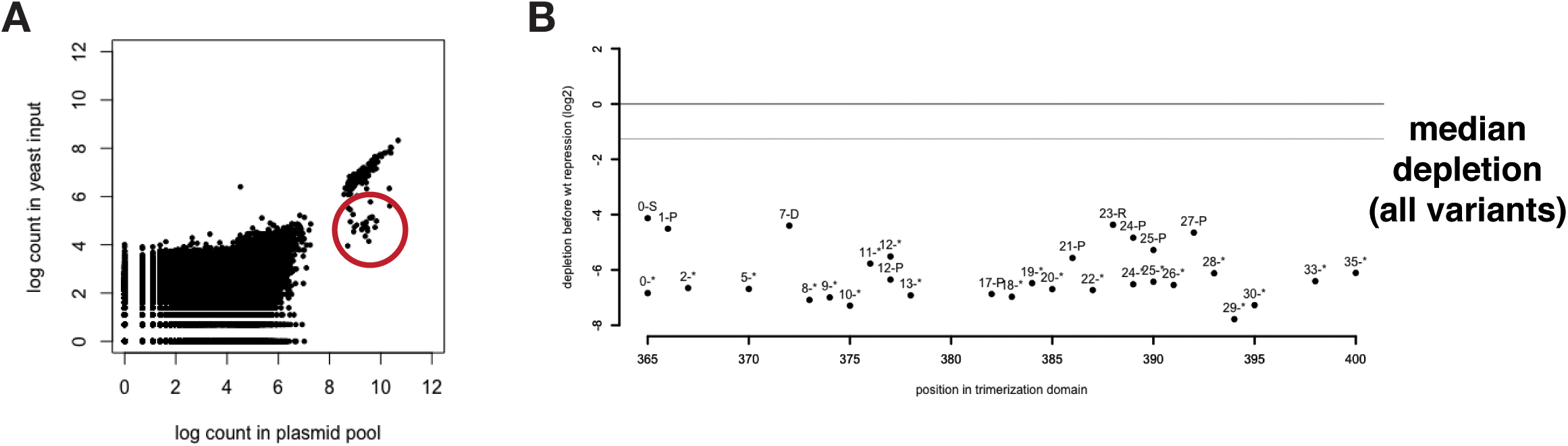
Hsf1 variants depleted in the population prior to repression of endogenous Hsf1 may represent dominant negative missense variants. (A) Scatterplot showing the frequency of each variant in the plasmid pool before yeast transformation (x-axis) and after yeast transformation and outgrowth without tet-induced repression of native *HSF1* (y-axis). Variants showing significant depletion (circled in red) while native *HSF1* is expressed presumably act through dominant negative inhibition of Hsf1. (B) Log2 fitness scores relative to wild type frequency during outgrowth of circled variants from (A) are plotted against their residue position in the Hsf1 trimerization domain. Gray horizontal line represents the median depletion of the rest of the library. The large majority of these variants represent nonsense mutations.

**Supplementary Figure 3.**
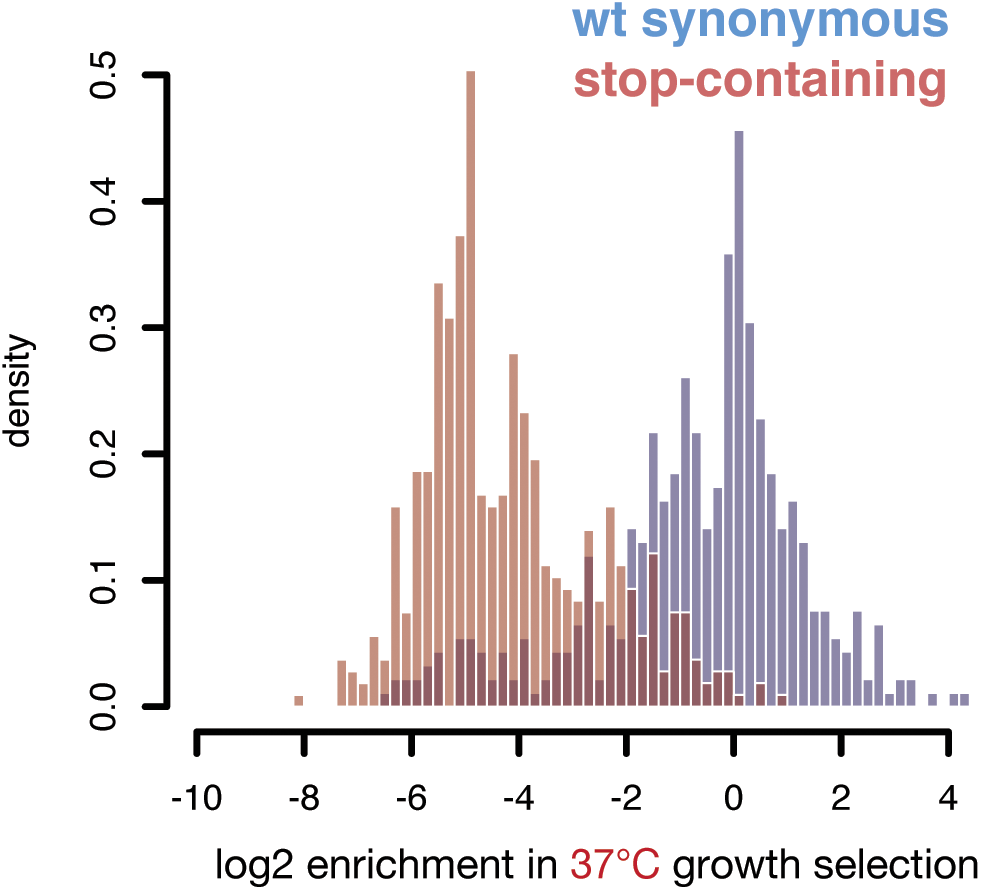
Synonymous Hsf1 variants show enrichment scores centered around zero. Histograms showing the log2 fitness scores of variants with synonymous codon changes to wild-type (blue) or variants containing stop codons (red) after growth at 37°C.

**Supplementary Figure 4.**
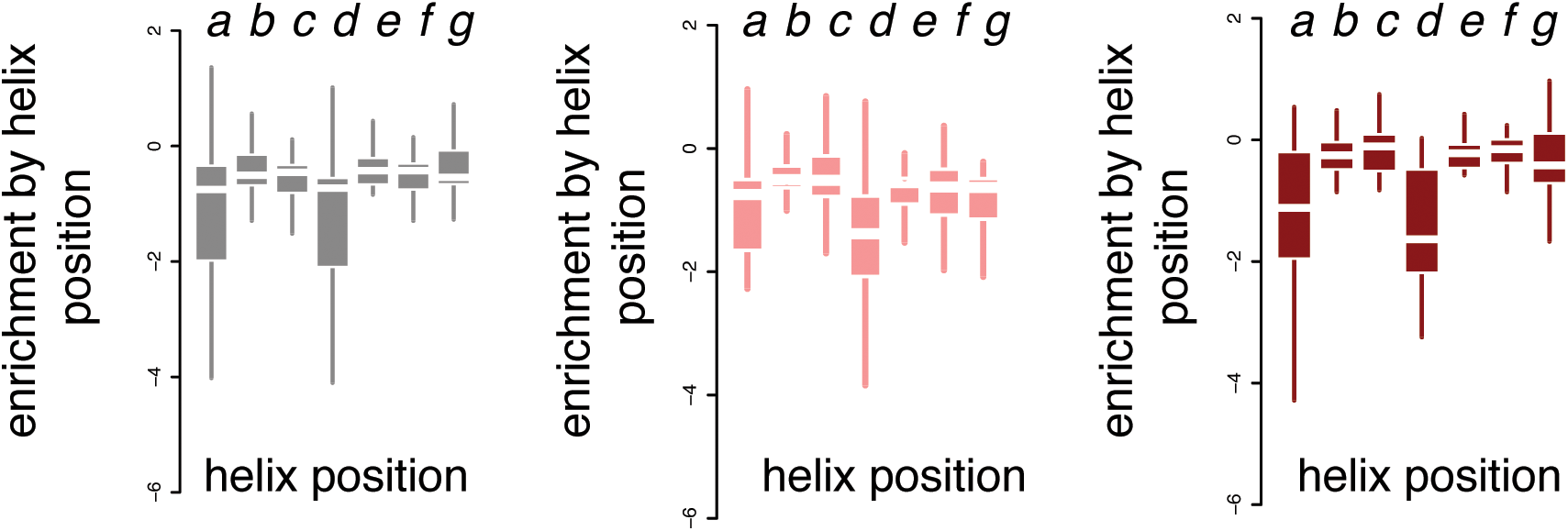
Small or charged substitutions do not have cumulatively negative effects outside of helix positions *a* and *d*. Boxplots showing log2 DMS enrichment scores of variants with substitutions to small or charged (K,R,D,E,S,G) residues at each site grouped by helix position (x-axis). Boxplot lines indicate median enrichment scores.

**Supplementary Figure 5.**
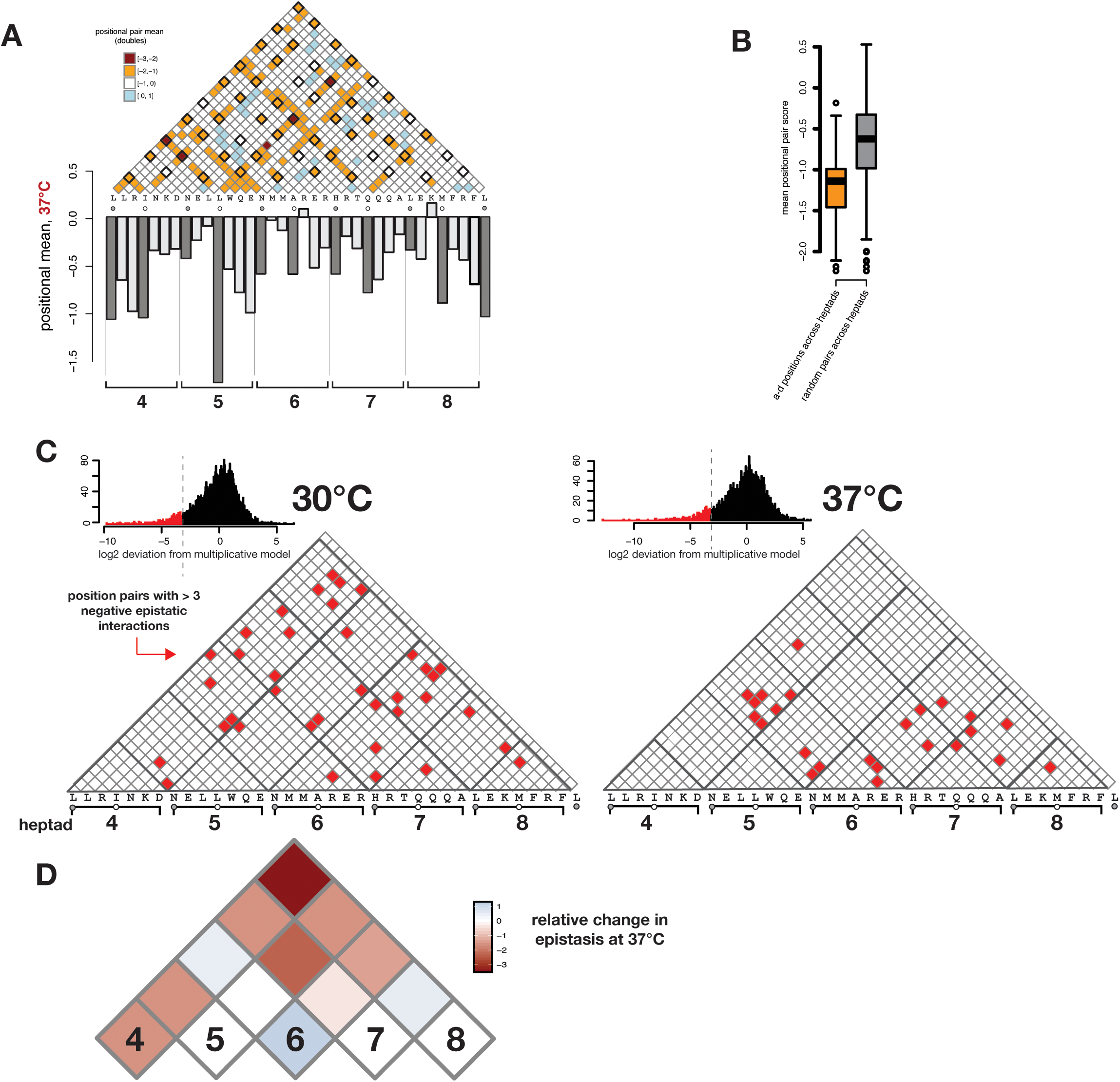
Analysis of double mutants supports the functional role of a and d positions and reveals patterns of epistatically-interacting sites underlying fewer long-distance contacts at 37°C. (A) Heatmap showing mean enrichment scores at 37°C of all variants found at each pair of sites. Darker shades of orange indicate site pairs for which the double mutants were more deleterious. Positional means generated for single amino acid substitutions at each residue (as in Fig. 3) are shown below for reference, with heptads indicated below. (B) Boxplot showing the mean enrichment scores of double mutants at a-d positions are more deleterious than pairs of mutations at other helix positions. (C) Pairs of sites are marked (red) where >3 double mutants show scores more negative than expected from multiplication of each constituent single (histogram shown above). The pattern of site pairs showing negative epistasis differs between 30°C (left) and 37°C (right). (D) The difference in epistasis scores between the two temperatures is presented as a heat map by heptad (4-8). Epistasis observed at distant heptads (e. g. between heptads 4 and 8) is reduced at 37°C compared to 30°C.

**Supplementary Figure 6.**
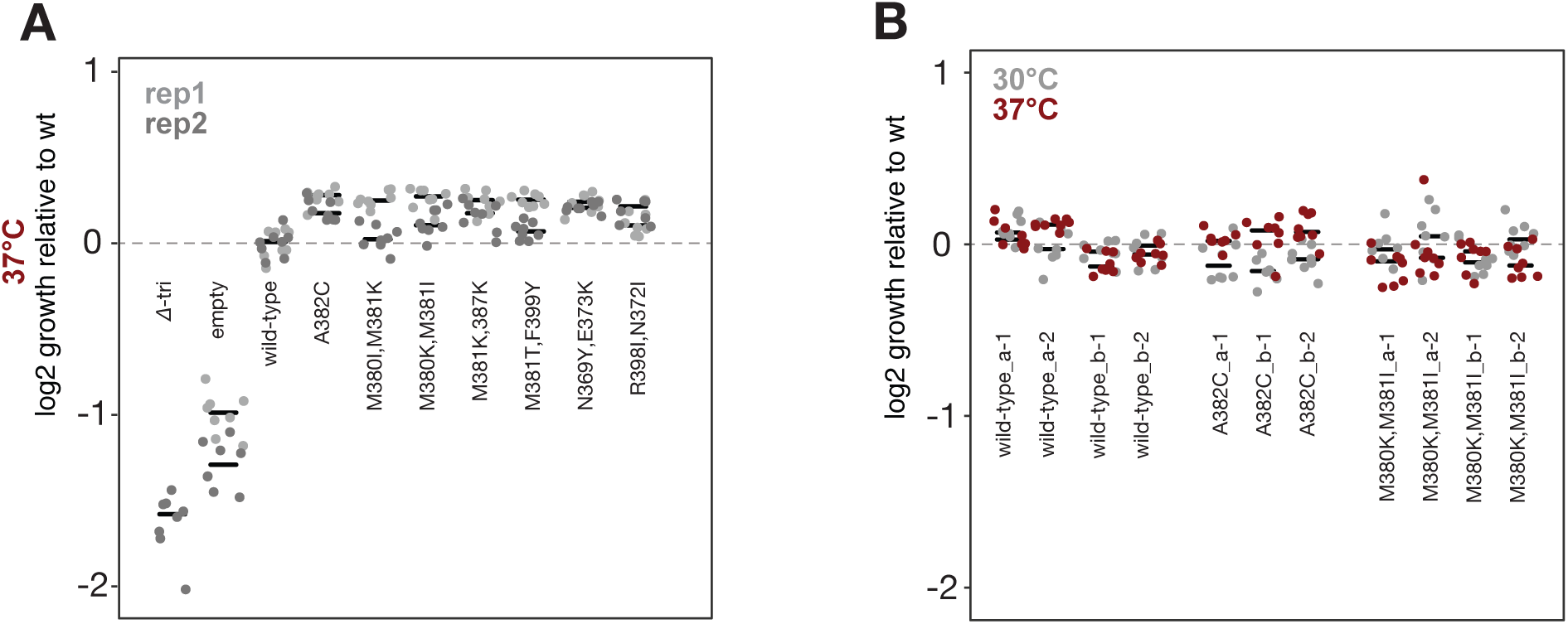
Growth rate validation of variants from large screen. (A) Seven *HSF1* variants identified in the DMS were independently validated in overnight plate reader growth experiments. *HSF1* variants were expressed in the conditions used during the DMS: endogenous *HSF1* was repressed with ATc and variant *HSF1* was expressed from a plasmid with the native HSF1 promoter. *HSF1* sequence contained the silent restriction enzyme sites used in DMS cloning. Dots represent technical replicate wells. Rep1 (light gray) and rep2 (dark gray) represent separate experiments performed with independent plasmid transformants. Data from rep2 is reproduced in Fig. 4C. The maximum slope of the growth curves is presented relative to the average rescue of a plasmid expressing wild-type *HSF1*. (B) *HSF1* variants were integrated into the chromosomal *HSF1* locus, under the native promoter and absent the silent restriction cloning sites. Clones from two different integration events were isolated for each genotype using *URA3* marker selection (a and b for each genotype) after which *URA3* was eliminated with counterselection (see Methods) to generate clones 1 and 2 of each integrated genotype. Overnight plate reader growth experiments were performed as above, at temperatures 30°C and 37°C. Individual dots represent technical replicate wells.

**Supplementary Figure 7.**
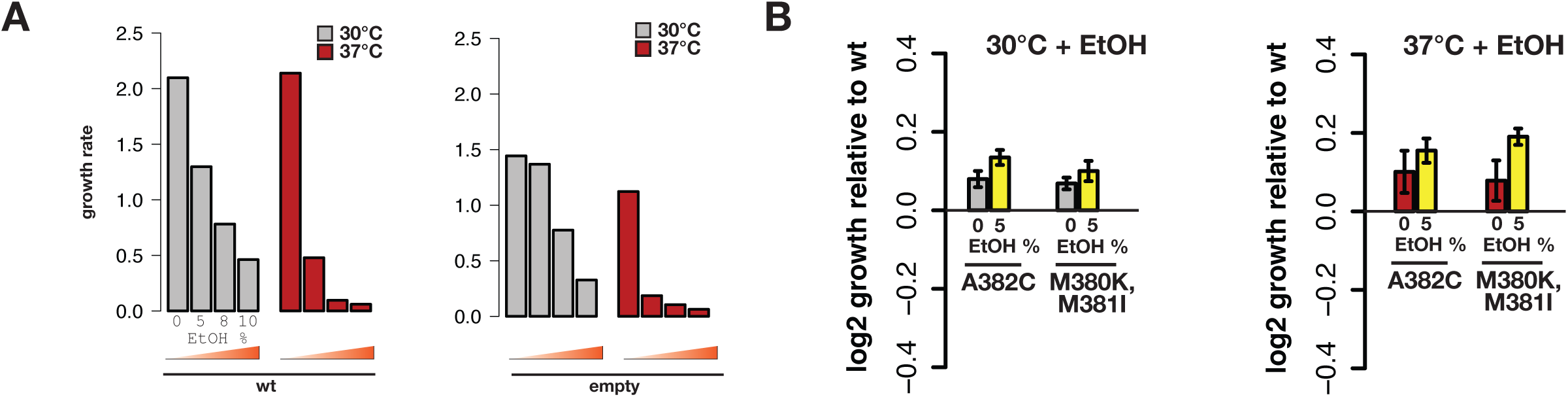
HSF1 variants improve growth under temperature and ethanol stress. (A) Yeast strains with endogenous HSF1 repressed and rescued with plasmid expression of either wild-type Hsf1 (wt) or an empty vector were grown in the presence of ethanol at 0%, 5%, 8%, or 10%. Overnight growth was assessed in a plate reader at 30°C or 37°C and the maximum growth velocity is presented. (B) Yeast strains were assessed as above but with expression of two different HSF1 variants: A382C and M380K,M381I, both of which were identified for their ability to confer enhanced growth at 37°C (Fig. 4C). The log2 of their maximum growth velocity relative to wild type is presented for growth in 0% or 5% ethanol at 30°C or 37°C. (Error bars represent standard error of the mean.)

**Supplementary Figure 8.**
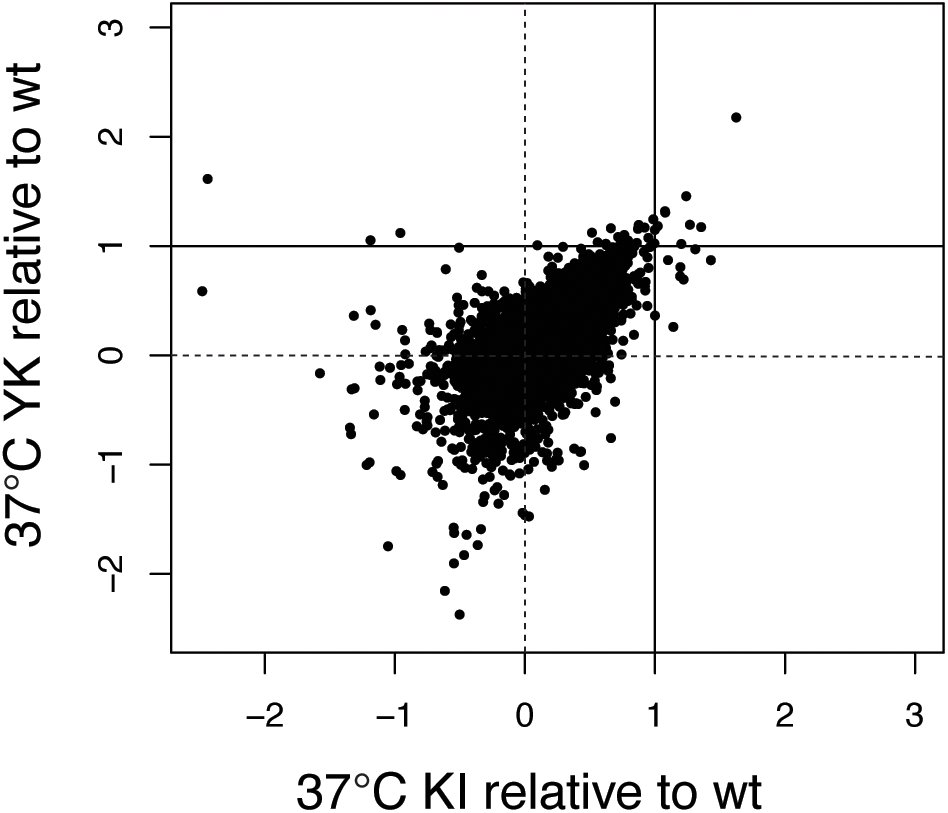
*HSF1* variants conferring enhanced growth phenotype showed similar patterns of upregulated genes under temperature stress. Scatterplot showing log2 relative expression level of each gene in M380K,M381I (KI) and N369Y,E373K (YK) variants at 37°C. Dotted guidelines represent no change in expression relative to wild type and solid guidelines indicate 2-fold changes in expression.

**Supplementary Figure 9.**
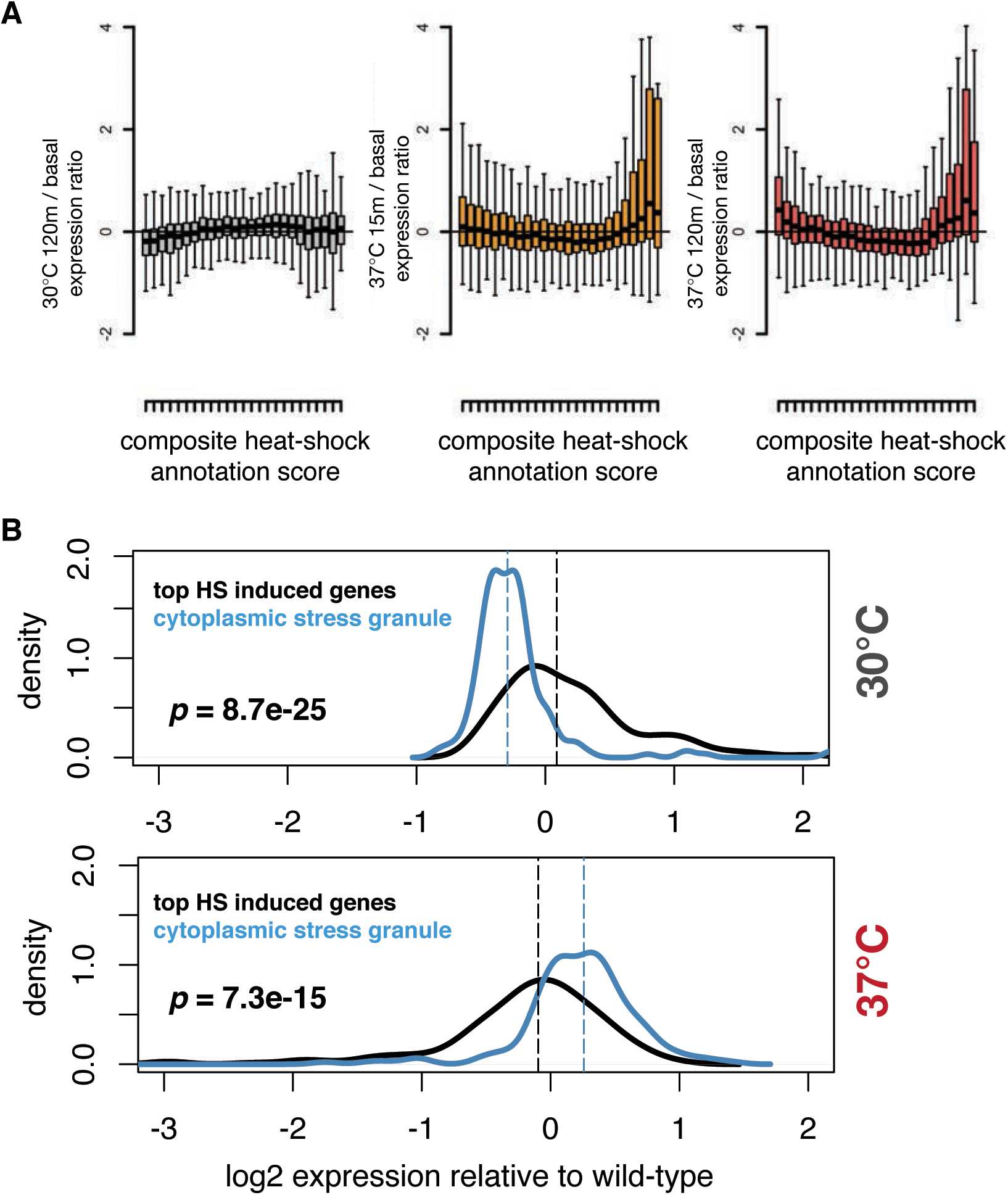
Yeast with expression of wild-type *HSF1* activate a core heat-shock response similar to previous data measuring gene expression in cells shifted to 37°C; variants show similar coreheat-shock response but alter expression of cytoplasmic stress granule-related genes at both temperatures. (A) Boxplots showing the RNA-seq expression levels of genes grouped by their composite rank among several expression studies in the Saccharomyces Genome Database testing 37°C induction. Each bin contains 200 genes, and the highest rank bin (further right on x-axis) includes the genes with the highest 37°C induction across all previous expression studies. For each bin, a boxplot of RNA-seq expression values are shown. The furthest left plot shows the RNA-seq expression values for cells harvested from the 30°C (non-stress) condition 120 minutes after repression of wild-type HSF1. In this case, bins containing frequently heat-induced genes show no significant expression increase. In contrast, the middle and right plots showing RNA-seq expression values from the 37°C (stress) condition 15 minutes and 120 minutes after repression of wild-type Hsf1, respectively, show increased expression of genes within bins containing the genes most frequently upregulated at 37°C in previous studies. (B) Density histograms showing the expression levels of two key sets of genes expressed in the two HSF1 variants (M380K,M381I and N369Y,E373K) relative to wild type. One set represents heat-shock induced genes from the top-ranked bin in (A) (black line) and the second represents all genes with the cytoplasmic stress granule annotation (blue line). While expression of cytoplasmic stress granule genes differs from wild type in the exceptional variants at both 30°C and 37°C, expression of the top heat-shock induced genes remains similar.

**Supplementary Figure 10.**
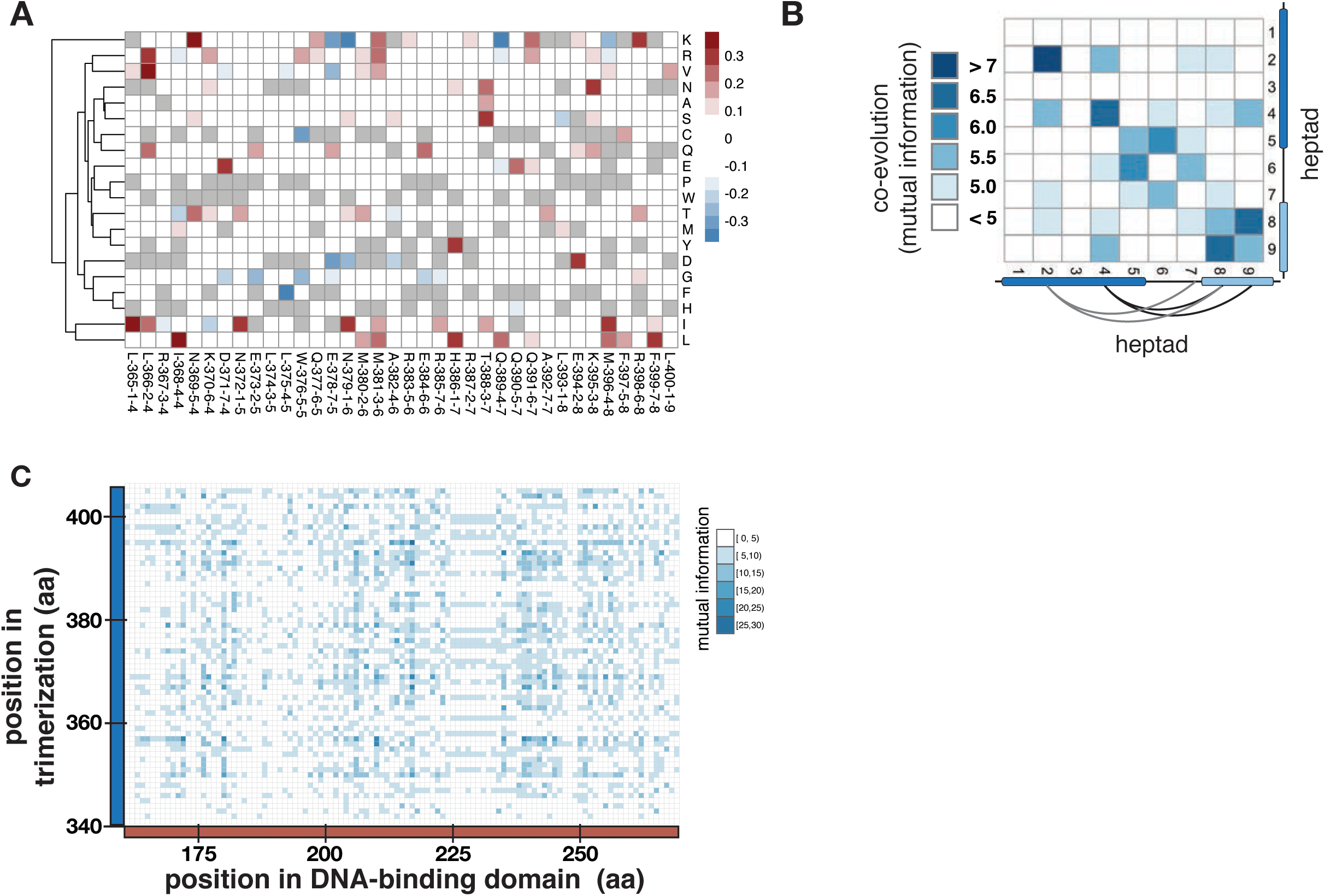
DMS data reveals particular amino acid substitutions enriched among the variants with greater fitness than wild type and analysis of fungal sequences reveals co-evolution across heptads in the oligomerization domain. (A) The pool of *HSF1* sequence variants that conferred improved growth at 37°C in the DMS was considered (regardless of how many mutations were present in each variant) and broken down by prevalence of any given substitution at each residue. A heatmap showing the frequency of each of these mutations compared to frequency in a random set of variants with near wild-type fitness is presented. Warmer shades indicate changes that are enriched in the exceptional variant population, while cooler shades identify changes that appear less frequently than expected in this group. (B) A mutual information heatmap of co-evolution of the HSF1 oligomerization domain in the ∼1200-species fungal database is presented, considered on the scale of heptads of the oligomerization domain. (C) Heatmap of mutual information between sites specifically in the DNA-binding domain of Hsf1 (red, x-axis) and the trimerization domain (blue, y-axis). Darker shades of blue in heatmap indicate higher levels of mutual information.

## References

1. Peteranderl, R. et al. Biochemical and biophysical characterization of the trimerization domain from the heat shock transcription factor. Biochemistry 38, 3559–69 (1999).

2. Sakurai, H. & Enoki, Y. Novel aspects of heat shock factors: DNA recognition, chromatin modulation and gene expression. FEBS J. 277, 4140–4149 (2010).

3. Hajdu-Cronin, Y. M., Chen, W. J. & Sternberg, P. W. The L-type cyclin CYL-1 and the heat-shock-factor HSF-1 are required for heat-shock-induced protein expression in Caenorhabditis elegans. Genetics 168, 1937–1949 (2004).

4. Morton, E. A. & Lamitina, T. Caenorhabditis elegans HSF-1 is an essential nuclear protein that forms stress granule-like structures following heat shock. Aging Cell 12, 112– 120 (2013).

5. Jedlicka, P., Mortin, M. A. & Wu, C. Multiple functions of Drosophila heat shock transcription factor in vivo. EMBO J. 16, 2452–2462 (1997).

6. Gallo, G. J., Prentice, H. & Kingston, R. E. Heat shock factor is required for growth at normal temperatures in the fission yeast Schizosaccharomyces pombe. Mol. Cell. Biol. 13, 749–761 (1993).

7. Sorger, P. K. & Pelham, H. R. B. Yeast Heat Shock Factor Is an Essential DNA-Binding Protein That Exhibits Temperature Dependent Phosphorylation. Cell 54, 855–864 (1988).

8. Xiao, X. Z. et al. HSF1 is required for extra-embryonic development, postnatal growth and protection during inflammatory responses in mice. EMBO J. 18, 5943–5952 (1999).

9. Metchat, A. et al. Mammalian heat shock factor 1 is essential for oocyte meiosis and directly regulates Hsp90α expression. J. Biol. Chem. 284, 9521–9528 (2009).

10. Hsu, A. L., Murphy, C. T. & Kenyon, C. Regulation of aging and age-related disease by DAF-16 and heat-shock factor. Science (80-.). 300, 1142–1145 (2003).

11. Dai, C., Whitesell, L., Rogers, A. B. & Lindquist, S. Heat Shock Factor 1 Is a Powerful Multifaceted Modifier of Carcinogenesis. Cell 130, 1005–1018 (2007).

12. Mendillo, M. L. et al. HSF1 drives a transcriptional program distinct from heat shock to support highly malignant human cancers. Cell 150, 549–562 (2012).

13. Scherz-Shouval, R. et al. The reprogramming of tumor stroma by HSF1 is a potent enabler of malignancy. Cell 158, 564–578 (2014).

14. Yin, J. et al. piR-823 contributes to colorectal tumorigenesis by enhancing the transcriptional activity of HSF1. Cancer Sci. 108, 1746–1756 (2017).

15. Singh, V. & Aballay, A. Heat-shock transcription factor (HSF)-1 pathway required for Caenorhabditis elegans immunity. Proc. Natl. Acad. Sci. U. S. A. 103, 13092–7 (2006).

16. Inouye, S. et al. Impaired IgG production in mice deficient for heat shock transcription factor 1. J. Biol. Chem. 279, 38701–38709 (2004).

17. Ma, X. et al. Celastrol protects against obesity and metabolic dysfunction through activation of a HSF1-PGC1α transcriptional axis. Cell Metab. 22, 695–708 (2015).

18. Minsky, N. & Roeder, R. G. Direct link between metabolic regulation and the heat-shock response through the transcriptional regulator PGC-1α. Proc. Natl. Acad. Sci. U. S. A. 112, E5669–E5678 (2015).

19. Jin, X., Qiao, A., Moskophidis, D. & Mivechi, N. F. Modulation of Heat Shock Factor 1 Activity through Silencing of Ser303/Ser307 Phosphorylation Supports a Metabolic Program Leading to Age-Related Obesity and Insulin Resistance. Mol. Cell. Biol. 38, (2018).

20. Qiao, A., Jin, X., Pang, J., Moskophidis, D. & Mivechi, N. F. The transcriptional regulator of the chaperone response HSF1 controls hepatic bioenergetics and protein homeostasis. J. Cell Biol. 216, 723–741 (2017).

21. Li, J., Labbadia, J. & Morimoto, R. I. Rethinking HSF1 in Stress, Development, and Organismal Health. Trends Cell Biol. 27, 895–905 (2017).

22. Douglas, P. M. et al. Heterotypic Signals from Neural HSF-1 Separate Thermotolerance from Longevity. Cell Rep. 12, 1196–1204 (2015).

23. Hahn, J., Hu, Z., Thiele, D. J. & Iyer, V. R. Genome-Wide Analysis of the Biology of Stress Responses through Heat Shock Transcription Factor. Mol. Cell. Biol. 24, 5249– 5256 (2004).

24. Pincus, D. et al. Genetic and epigenetic determinants establish a continuum of Hsf1 occupancy and activity across the yeast genome. Mol. Biol. Cell 29, 3168–3182 (2018).

25. Clos, J., Rabindran, S., Wisniewski, J. & Wu, C. Induction temperature of human heat shock factor is reprogrammed in a Drosophila cell environment. Nature 364, 252–5 (1993).

26. Hentze, N., Breton, L. Le, Wiesner, J., Kempf, G. & Mayer, M. P. Molecular mechanism of thermosensory function of human heat shock transcription factor Hsf1. eLife 5, e11576 (2016).

27. Soncin, F. et al. Transcriptional activity and DNA binding of heat shock factor-1 involve phosphorylation on threonine 142 by CK2. Biochem. Biophys. Res. Commun. 303, 700– 706 (2003).

28. Hashikawa, N., Yamamoto, N. & Sakurai, H. Different mechanisms are involved in the transcriptional activation by yeast heat shock transcription factor through two different types of heat shock elements. J. Biol. Chem. 282, 10333–10340 (2007).

29. Satyal, S. H., Chen, D., Fox, S. G., Kramer, J. M. & Morimoto, R. I. Negative regulation of the heat shock transcriptional response by HSBP1. Genes Dev. 12, 1962–1974 (1998).

30. Tan, K. et al. Mitochondrial SSBP1 protects cells from proteotoxic stresses by potentiating stress-induced HSF1 transcriptional activity. Nat. Commun. 6, 1–15 (2015).

31. Neef, D. W., Jaeger, A. M. & Thiele, D. J. Genetic selection for constitutively trimerized human hsf1 mutants identifies a role for coiled-coil motifs in DNA binding. G3 Genes, Genomes, Genet. 3, 1315–1324 (2013).

32. Takemori, Y. et al. Mutational analysis of human heat-shock transcription factor 1 reveals a regulatory role for oligomerization in DNA-binding specificity. Biochem. J. 424, 253– 261 (2009).

33. Sorger, P. K. & Nelson, H. C. M. Trimerization of a yeast transcriptional activator via a coiled-coil motif. Cell 59, 807–813 (1989).

34. Zuo, J., Baler, R., Dahl, G. & Voellmy, R. Activation of the DNA-binding ability of human heat shock transcription factor 1 may involve the transition from an intramolecular to an intermolecular triple-stranded coiled-coil structure. Mol. Cell. Biol. 14, 7557–7568 (1994).

35. Giardina, C. & Lis, J. T. Dynamic protein-DNA architecture of a yeast heat shock promoter. Mol. Cell. Biol. 15, 2737–2744 (1995).

36. Crick, F. H. C. The packing of α-helices: simple coiled-coils. Acta Crystallogr. 6, 689– 697 (1953).

37. McLachlan, A. D. & Stewart, M. Tropomyosin coiled-coil interactions: Evidence for an unstaggered structure. J. Mol. Biol. 98, 293–304 (1975).

38. Clos, J. et al. Molecular cloning and expression of a hexameric Drosophila heat shock factor subject to negative regulation. Cell 63, 1085–1097 (1990).

39. Peteranderl, R. & Nelson, H. C. M. Trimerization of the Heat Shock Transcription Factor by a Triple-Stranded α-Helical Coiled-Coil. Biochemistry 31, 12272–12276 (1992).

40. Xu, W. et al. The influence of the mating type on virulence of Mucor irregularis. Sci. Rep. 7, 1–12 (2017).

41. Akaza, N. et al. Malassezia globosa tends to grow actively in summer conditions more than other cutaneous Malassezia species. J. Dermatol. 39, 613–616 (2012).

42. Vishniac, H. S. & Hempfling, W. P. Cryptococcus vishniacii sp. nov., an Antarctic Yeast. Int. J. Syst. Bacteriol. 29, 153–158 (1979).

43. van den Brink, J., Facun, K., de Vries, M. & Stielow, J. B. Thermophilic growth and enzymatic thermostability are polyphyletic traits within Chaetomiaceae. Fungal Biol. 119, 1255–1266 (2015).

44. Wang, H., Mayhew, D., Chen, X., Johnston, M. & Mitra, R. D. Calling Cards enable multiplexed identification of the genomic targets of DNA-binding proteins. Genome Res. 21, 748–755 (2011).

45. Zhou, W., Dorrity, M. W., Bubb, K. L., Queitsch, C. & Fields, S. Binding and regulation of transcription by yeast Ste12 variants to drive mating and invasion phenotypes. Genetics 214, 397–407 (2020).

46. Yamamoto, A., Mizukami, Y. & Sakurai, H. Identification of a novel class of target genes and a novel type of binding sequence of heat shock transcription factor in Saccharomyces cerevisiae. J. Biol. Chem. 280, 11911–11919 (2005).

47. Palii, C. G. et al. Differential genomic targeting of the transcription factor TAL1 in alternate haematopoietic lineages. EMBO J. 30, 494–509 (2011).

48. Li, C. et al. Genome-wide characterization of cis-acting DNA targets reveals the transcriptional regulatory framework of Opaque2 in maize. Plant Cell 27, 532–545 (2015).

49. Wang, S. et al. Target analysis by integration of transcriptome and ChIP-seq data with BETA. Nat. Protoc. 8, 2502 (2013).

50. Opulente, D. A. et al. Factors driving metabolic diversity in the budding yeast subphylum. BMC Biol. 16, 1–15 (2018).

51. Kurtzman, C., Fell, J. W. & Boekhout, T. The yeasts: a taxonomic study. (Elsevier, 2011).

52. Nover, L. et al. Arabidopsis and the heat stress transcription factor world: How many heat stress transcription factors do we need? Cell Stress Chaperones 6, 177–189 (2001).

53. Chen, Y., Barlev, N. A, Westergaard, O. & Jakobsen, B. K. Identification of the C-terminal activator domain in yeast heat shock factor: independent control of transient and sustained transcriptional activity. EMBO J. 12, 5007–18 (1993).

54. Jakobsen, B. K. & Pelham, H. R. A conserved heptapeptide restrains the activity of the yeast heat shock transcription factor. EMBO J. 10, 369–375 (1991).

55. Puustinen, M. C. & Sistonen, L. Molecular Mechanisms of Heat Shock Factors in Cancer. Cells 9, 1–20 (2020).

56. Hashikawa, N., Mizukami, Y., Imazu, H. & Sakurai, H. Mutated yeast heat shock transcription factor activates transcription independently of hyperphosphorylation. J. Biol. Chem. 281, 3936–3942 (2006).

57. Jolma, A. et al. DNA-dependent formation of transcription factor pairs alters their binding specificity. Nature 527, 384–388 (2015).

58. Rohs, R., West, S. M., Sosinsky, A., Liu, P. & Mann, R. S. The role of DNA shape in protein-DNA recognition. Nature 461, (2009).

59. Ali, M. H. & Imperiali, B. Protein oligomerization: How and why. Bioorganic Med. Chem. 13, 5013–5020 (2005).

60. Ulmasov, T., Hagen, G. & Guilfoyle, T. J. Dimerization and DNA binding of auxin response factors. Plant J. 19, 309–319 (1999).

61. Dorrity, M. W., Cuperus, J. T., Carlisle, J. A., Fields, S. & Queitsch, C. Preferences in a trait decision determined by transcription factor variants. Proc. Natl. Acad. Sci. 115, E7997–E8006 (2018).

62. Mumberg, D., Müller, R. & Funk, M. Yeast vectors for the controlled expression of heterologous proteins in different genetic backgrounds. Gene 156, 119–22 (1995).

63. Mnaimneh, S. et al. Exploration of essential gene functions via titratable promoter alleles. Cell 118, 31–44 (2004).

64. Edgar, R. C. MUSCLE: Multiple sequence alignment with high accuracy and high throughput. Nucleic Acids Res. 32, 1792–1797 (2004).

65. Crooks, G. E., Hon, G., Chandonia, J.-M. & Brenner, S. E. WebLogo: a sequence logo generator. Genome Res. 14, 1188–1190 (2004).

66. Simonetti, F. L., Teppa, E., Chernomoretz, A., Nielsen, M. & Marino Buslje, C. MISTIC: mutual information server to infer coevolution. Nucleic Acids Res. 41, W8--W14 (2013).

67. Gietz, R. D. & Woods, R. A. Transformation of yeast by lithium acetate/single-stranded carrier DNA/polyethylene glycol method. Methods Enzymol. 350, 87–96 (2002).

68. Hoffman, C. S. & Winston, F. A ten-minute DNA preparation from yeast efficiently releases autonomous plasmids for transformaion of Escherichia coli. Gene 57, 267–272 (1987).

69. Bailey, T. L., Johnson, J., Grant, C. E. & Noble, W. S. The MEME suite. Nucleic Acids Res. 43, W39--W49 (2015).

70. Teixeira, M. C. et al. YEASTRACT: an upgraded database for the analysis of transcription regulatory networks in Saccharomyces cerevisiae. Nucleic Acids Res. 46, D348--D353 (2018).

71. Reimand, J. et al. g: Profiler—a web server for functional interpretation of gene lists (2016 update). Nucleic Acids Res. 44, W83--W89 (2016).

72. Cuperus, J. T., Lo, R. S., Shumaker, L., Proctor, J. & Fields, S. A tetO Toolkit to Alter Expression of Genes in Saccharomyces cerevisiae. ACS Synth. Biol. 4, 842–852 (2015).

73. Supek, F., Bošnjak, M., Škunca, N. & Šmuc, T. Revigo summarizes and visualizes long lists of gene ontology terms. PLoS One 6, (2011).

74. Storici, F. & Resnick, M. A. The delitto perfetto approach to in vivo site-directed mutagenesis and chromosome rearrangements with synthetic oligonucleotides in yeast. Methods Enzymol. 409, 329–345 (2006).

75. Åkerfelt, M., Morimoto, R. I. & Sistonen, L. Heat shock factors: Integrators of cell stress, development and lifespan. Nat. Rev. Mol. Cell Biol. 11, 545–555 (2010).

76. Littlefield, O. & Nelson, H. C. M. A new use for the’wing’of the’winged’helix-turn-helix motif in the HSF–DNA cocrystal. Nat. Struct. Mol. Biol. 6, 464–470 (1999).

77. Cicero, M. P. et al. The wing in yeast heat shock transcription factor (HSF) DNA-binding domain is required for full activity. Nucleic Acids Res. 29, 1715–1723 (2001).

